# Developmental and evolutionary constraints on olfactory circuit selection

**DOI:** 10.1101/2020.12.22.423799

**Authors:** Naoki Hiratani, Peter E. Latham

## Abstract

Across species, neural circuits show remarkable regularity, suggesting that their structure has been driven by underlying optimality principles. Here, we ask whether we can predict the neural circuitry of diverse species by optimizing the neural architecture to make learning as efficient as possible. We focus on the olfactory system, primarily because it has a relatively simple evolutionarily conserved structure, and because its input and intermediate layer sizes exhibits a tight allometric scaling. In mammals, it has been shown that the number of neurons in layer 2 of piriform cortex scales as the number of glomeruli (the input units) to the 3/2 power; in invertebrates, we show that the number of mushroom body Kenyon cells scales as the number of glomeruli to the 7/2 power. To understand these scaling laws, we model the olfactory system as a three layered nonlinear neural network, and analytically optimize the intermediate layer size for efficient learning from a limited number of samples. We find that the 3/2 scaling observed in mammals emerges naturally, both in full batch optimization and under stochastic gradient learning. We extended the framework to the case where a fraction of the olfactory circuit is genetically specified, not learned. We show numerically that this makes the scaling law steeper when the number of glomeruli is small, and we are able to recover the 7/2 scaling law observed in invertebrates. This study paves the way for a deeper understanding of the organization of brain circuits from an evolutionary perspective.

## I. INTRODUCTION

Brains exhibit a large range of cell types, connectivity patterns, and organizational structures, at both micro and macro scales. There is a rich history in neuroscience of explaining these structures from a normative point of view [1–3]. Most of that work focused on computation, in the sense that it asked what circuit, and connection strengths, lead to optimal performance on a particular task. However, the connection strengths have to be learned, and model selection theory tells us that the efficiency of learning depends crucially on architecture, especially when a limited number of trials is available [4–8]. In this study, we attempt to understand the organizational structure of the brain from a model selection perspective, hypothesizing that evolution optimized the brain for efficiency of learning.

Here we focus on the olfactory system, primarily because it has a relatively simple, evolutionarily conserved, predominantly feedforward structure [9–11]. In particular, odorants are first detected by olfactory sensory neurons; from there, olfactory information is transmitted to glomeruli. The number of glomeruli, however, varies widely across species, from between 10 and 100 in insects to ∼1000 in mammals. The question we address here is: how does the number of glomeruli affect downstream circuitry? And in particular, what downstream circuitry would best help the animal survive? The tradeoffs that go into answering this question are in principle straightforward: more complicated circuitry (i.e., more parameters) can do a better job accurately predicting reward and punishment, but, because there are more parameters, there is a danger of overfitting [4, 7, 8]. And even if learning is performed with sample-by-sample updates to avoid overfitting, learning tends to be slower in complicated circuitry, as typically more samples are required [12, 13]. Navigating these tradeoffs for a given architecture is reasonably straightforward. The architecture, though, must come from biology. For that we take inspiration from the olfactory system of both mammals and invertebrates.

In the mammalian olfactory system, information from the glomeruli is transmitted to mitral/tufted cells, then to layer 2 of piriform cortex among others, and then mainly to layer 3; after that, information is passed on to higher order cortical areas [9, 11]. Thus, although many studies suggest that reciprocal interactions between mitral/tufted cells and granule cells are also important for olfactory processing [14–16], as a first-order approximation the olfactory system can be modeled as a feedforward neural network. Moreover, because sister mitral cells receiving input from the same glomeruli are highly correlated, both with each other and with the glomeruli from which they receive input [17], the olfactory network essentially has three layers: an input layer corresponding to glomeruli, a hidden layer corresponding to layer 2 of piriform cortex, and an output layer corresponding to layer 3.

Based on this picture, in our analysis we use an architecture corresponding to a three layer feedforward network. The size of the input layer is the number of glomeruli, and we assume that each unit of the output layer is extracting a different feature of the olfactory input, such as expected reward or punishment, or a behaviorally relevant concept. Consequently, we focus on the hidden layer. For that we ask: how many units should the hidden layer have? That question was chosen partly because its answer provides insight into learning principles in general, and partly because it was recently addressed experimentally: Srinivasan and Stevens found, based on six mammalian species, a very tight relationship between the number of glomeruli and the number of neurons in layer two of piriform cortex (Fig. 1A; data taken from [18]). More precisely, using *L*_*x*_ to denote the input layer size (the number of glomeruli), and *L*_*h*_ to denote the hidden layer size (the number of neurons in layer 2 of piriform cortex), they found the approximate scaling law 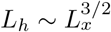.

**FIG. 1.**
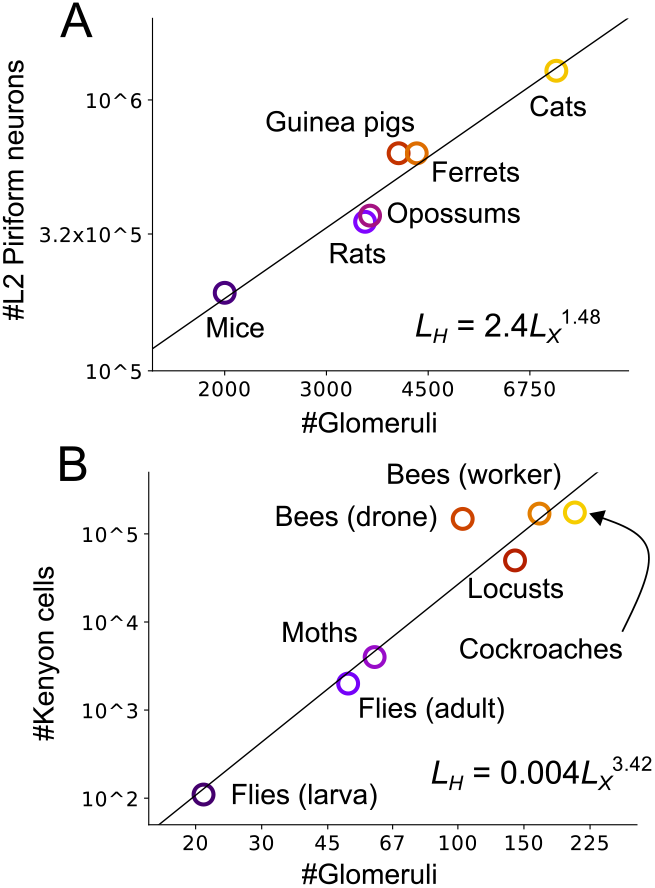
**A)** Scaling law in mammalian olfactory circuits. Data was replicated from supplementary tables S2 and S3 of Srinivasan & Stevens, 2019. **B)** Scaling law in invertebrate olfactory circuits. See SI §1.1 for the details.

Motivated by this result, we asked whether a similar scaling law holds for the invertebrate olfactory system. Like their mammalian counterparts, odors detected by olfactory sensory neurons converge to glomeruli. After that, though, the circuitry differs. Glomeruli send information to the projection neurons [11], which mainly extend synapses onto mushroom body Kenyon cells and lateral horn neurons [19]. The latter is mostly related to innate olfactory processing [20], so we focus on the mushroom body, which transmits information to higher-order regions through mushroom body output neurons. Thus, the invertebrate olfactory system can also be modeled as a three layer neural network: an input layer corresponding to glomeruli, a hidden layer corresponding to Kenyon cells, and an output layer corresponding to mushroom body output neurons [3, 21].

A literature survey of the number of glomeruli and Kenyon cells of various insects [22–34] (see SI §1.1 for details) yielded a scaling law, as in the mammalian olfactory system, but with an exponent of about 7/2 rather than 3/2 (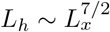, as shown in Fig. 1B). Drone (male) bees are the clear outlier, but that is reasonable considering the caste system of honey bees that puts the drones under unique ecological pressure; for instance, the drones are the only ones among the seven insects listed that do not engage in foraging. It should be noted that the data was not properly controlled, as it was collected from different sources, and in some cases in different eras. Moreover, for the locust, we used the number of olfactory receptor genes instead of the number of glomeruli; that is because their micro-glomeruli structure makes direct comparison with other species difficult [35]. In addition, the mushroom body also takes part in visual processing in bees and cockroaches [36].

Several normative hypotheses have been offered to explain the population size of sensory circuits. One line of theoretical work showed that expansion in the hidden layer is beneficial for sensory coding [3, 37, 38], but it remains elusive how much expansion is optimal, because in these studies, more expansion was in principle always better. Other studies estimated the optimal population size in multiple layers from a width-depth tradeoff, assuming that the total number of neurons is fixed [39, 40] by external factors such as a constraint on energy [41]. However, this energy constraint should be violated if increasing the number of neurons improves foraging ability, resulting in a better energy budget [42]. Evaluation of the optimal population size was also attempted from other biological constraints such as synaptic [43] and neuronal [44] noise. While these models provided insight into circuit structure, none were able to provide a quantitative explanation for the population sizes of circuits across different species. Srinivasan and Stevens, on the other hand, offered several explanations, based on coding efficiency and geometry, for the scaling in mammals [18]. While those explanations are reasonable candidate hypotheses, they are more abstract than mechanistic, and do not explain the scaling seen in invertebrates.

Here we develop a mechanistic explanation of the scaling laws, focusing on the fact that the transformation from glomeruli to piriform cortex (for mammals), or from glomeruli to mushroom body output neurons (for invertebrates), has to be learned from a limited number of samples. For that we apply model selection theory, in which the primary constraint on the circuit is the poverty of the teaching signals and resultant overfitting [4, 7, 8]. The olfactory circuit has to tune its numerous synaptic weights from very sporadic, low-dimensional reward signals in the natural environment [45, 46], so this constraint should be highly relevant. Therefore, we formulate the problem of olfactory circuit design as a model selection problem, then analytically derive the optimal hidden layer size under various learning rules and nonlinearities.

Not surprisingly (because learning takes time) we find that the optimal hidden layer size depends on the lifetime of the organism. Using observed lifetimes, we recover the 3/2 scaling found in mammals. However, our theory cannot capture the 7/2 power law found in invertebrates. That is because traditional model selection theory fails to take into account the fact that neural circuits are at least partially genetically specified. In particular, rich innate connectivity structure is known to exist in the invertebrate olfactory systems [20, 47]. Thus, we extend the framework to the case where a fixed genetic budget can be used to specify connections, and consider how that affects scaling. The budget we used – about 2000 bits – had little effect on the scaling of the mammalian circuit, primarily because mammals have a large number of glomeruli, for which a complicated downstream circuit is needed to achieve good performance – far more complicated than could be constructed by 2000 bits. However, it had a large effect on invertebrates, which contain far fewer glomeruli. Using this extended framework, we were able to recover the observed 7/2 power law without disrupting the 3/2 power law found in mammals. These results shed light on potential constraints on the development and evolution of neural circuitry.

## II. RESULTS

To determine scaling in the olfactory system, we use a teacher-student framework [13, 48, 49]: we postulate a teacher network, which reflects the true mapping from odors to reward or punishment in the environment, and model the circuit in the animal’s olfactory system using the same overall architecture, but with different nonlinearities and a different number of neurons in the hidden layer (see Fig. 2). We determine the optimal hidden layer size under several scenarios: batch learning and stochastic gradient learning, and with and without information about the weights supplied by the genome.

**FIG. 2.**
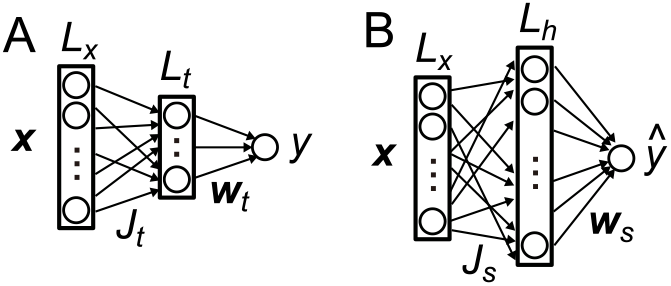
Network models. **A**) Olfactory environment (teacher). **B**) Olfactory circuit that models the environment (student).

### A. The model

Let us denote the olfactory input at the level of glomeruli as 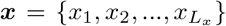, and the corresponding reward, or punishment, as *y*. We consider a student-teacher model, and define the true relationship between ***x*** and *y* in the environment by a three layer “teacher” network (Fig. 2A),

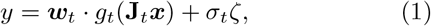

where *g*_*t*_ is a pointwise nonlinear activation function of the hidden neurons, and *ζ* is Gaussian noise, added because the relationship between input and reward is stochastic in real world situations. Throughout the text we use bold capital letters to denote matrices and bold small letter for vectors. Vectors are defined as column vectors, a superscript *T* denotes transpose (indicating a row vector), and for readability we use a dot product to denote the inner product between two vectors. We sampled **J**_*t*_, ***w***_*t*_, and ***x*** from independent Gaussian distributions for analytical tractability.

As discussed above, we model the olfactory circuits of both vertebrates and invertebrates as a three layer neural network (Fig. 2B),

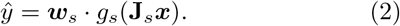

For simplicity, we assume that **J**_*s*_ is fixed and random, with elements drawn from an independent Gaussian distribution. Only the readout weights, ***w***_*s*_, are learned from data. This is a good approximation for the invertebrate olfactory system, as the connection from the projection neurons to Kenyon cells are indeed mostly random [25] and fixed [50]. In the mammalian system, the connection from mitral/tufted cells to piriform cortex, which corresponds to **J**_*s*_, is suggested to be plastic [51]. However, it is thought that those connections are mainly shaped by unsupervised learning, but are seldom modulated by reward, as odor representation in layer 2 of piriform cortex is relatively stable under reward-based learning [52, 53].

The objective of learning is to predict the true reward signal, *y*, given the input, ***x***. Using the mean squared error as the loss, the generalization error is written

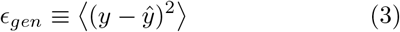

where here and in what follows angle brackets indicate an average over the input, ***x***, and the teacher noise, *ζ*. Under this problem setting, we ask what hidden layer size, *L*_*h*_, minimizes the generalization error, *∈*_*gen*_, when ***w***_*s*_ is learned from *N* training samples. In particular, we investigate how the optimal hidden layer size scales with the input layer size, *L*_*x*_. Intuitively, when the hidden layer size is too small, the neural network is not expressive enough, so the generalization error tends to be large even after an infinite number of training samples. On the other hand, if the hidden layer is too large relative to the number of training samples, the network becomes prone to over-fitting, again resulting in poor generalization. Here we solve this model selection problem analytically.

### B. Generalization error

When the learning rule is unbiased, the generalization error consists of two components: the approximation error, which arises because we do not have a perfect model (we use **J**_*s*_ rather than the true weight, **J**_*t*_, to model the output, *y*, and we may have a different nonlinearity and hidden layer size), and the estimation error, which arises because we use a finite number of training samples [6– 8]. Inserting Eqs. (1) and (2) into (3), we can write the generalization error in terms of these two components,

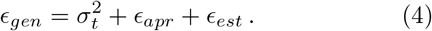

The approximation error, *∈*_*apr*_ (the error under the optimal weight 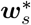), is given by

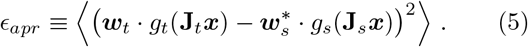

The estimation error, *∈*_*est*_ (the error induced by using the learned weight, ***w***_*s*_, rather than the optimal one, 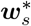), is

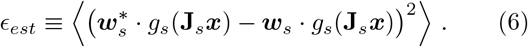

Note that under an appropriate learning rule, *ϵ*_*est*_ converges to zero in the limit of an infinite number of training samples (*N* → ∞).

We focus first on the approximation error, *ϵ*_*apr*_, which depends on the optimal weight, 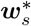. That weight is found by minimizing ⟨ (*y* − *ŷ*)^2^⟩ with respect to ***w***_*s*_, with *y* and *ŷ* given in Eqs. (1) and (2), respectively. This is a linear regression problem, and so 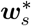 is given by the usual expression,

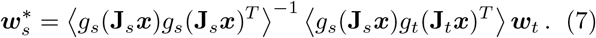

To compute 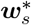, we need to invert a matrix. That is nontrivial because *g*_*s*_() is a nonlinear function and the components of **J**_*s*_***x*** are correlated,

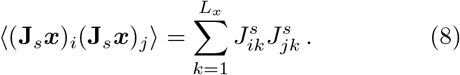

Because the 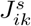 are independent random variables, the off-diagonal elements are smaller than the diagonal elements by a factor of *L*_*x*_. We can, therefore, compute 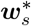 as an expansion in powers of 1*/L*_*x*_, multiplied by *L*_*h*_ (because there are factor of *L*_*h*_ more off-diagonal than diagonal elements). Working to second order in 1*/L*_*x*_, we show in SI §3 that

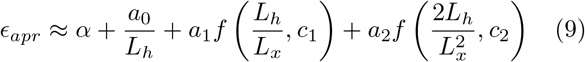

where

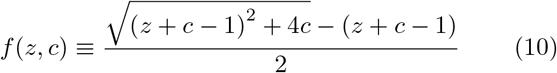

is a monotonically decreasing function of *z*: *f* (0, *c*) = 1 and *f* (*z, c*) → *c/z* when *z* » 1. All constants are *𝒪* (1); their values depend only on the nonlinearities *g*_*s*_() and *g*_*t*_(). Note that *ϵ*_*apr*_ does not explicitly depend on the teacher network size *L*_*t*_. That holds so long as *L*_*t*_ » 1 (see SI, Eqs. (36)-(39)).

As shown in Fig. 3 (blue line), *ϵ*_*apr*_ is a monotonically decreasing function of *L*_*h*_. That function derives its shape from the three *L*_*h*_-dependent terms in Eq. (9) (excluding *α*, which is a small constant): the second term, *α*_0_*/L*_*h*_, decays to zero when *L*_*h*_ is large compared to 1, the third decays to zero when *L*_*h*_ is large compared to *L*_*x*_, and the last decays to zero when *L*_*h*_ is large compared to 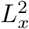. Essentially, as *L*_*h*_ increases, the effect of the off-diagonal elements of the covariance matrix in Eq. (7) increase, and the model becomes effectively more expressive (and thus lowers the approximation error). Although a number of approximations were made in deriving Eq. (9), the theoretical prediction matches well the numerical simulations (points in Fig. 3) for a wide range of *L*_*h*_.

**FIG. 3.**
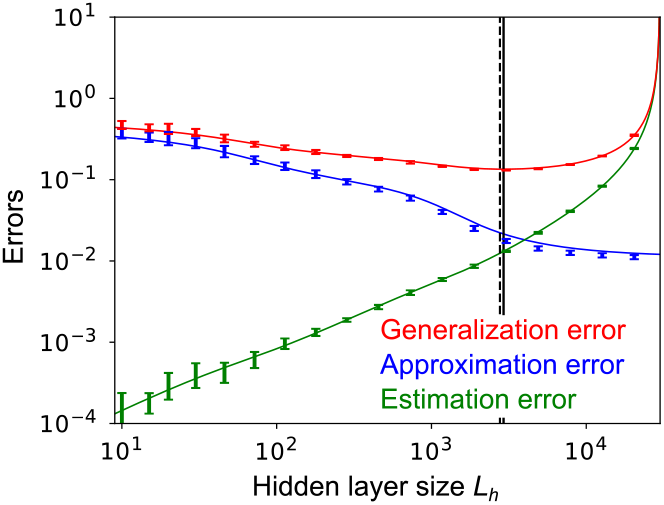
Generalization, approximation, and estimation error at *L*_*x*_ = 50, *N* = 30000, under various hidden layer sizes *L*_*h*_. Lines are analytical results (red, generalization error, Eq. (12); blue, approximation error, Eq. (9), and green, estimation error, Eq. (11)); points are from numerical simulations (see SI §7.3 for details). Here, and in all figures except Figs. 4D and E, both *g*_*t*_ and *g*_*s*_ are rectified linear functions (*g*_*t*_(*u*) = *g*_*s*_(*u*) = max(0, *u*)). In all figures except Fig. 6 we use 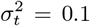 for the noise in the teacher circuit, and in all figures the hidden layer size of the teacher network is fixed at *L*_*t*_ = 500. Error-bars represent the standard deviation over 10 simulation trials.

To complete the picture of the generalization error, we need the estimation error – the error associated with finite training data. For that it matters how we learn ***w***_*s*_. There are two main choices: maximum likelihood estimation (MLE) and stochastic gradient descent (SGD). We consider MLE learning first. Although it is not biologically plausible (it requires the learner to compute, and invert, a covariance matrix after seeing all the data), we consider it first because it is reasonably straightforward. After that, we consider the more realistic case of SGD. Both exhibit the 3/2 scaling found in the mammalian olfactory circuit.

In SI §4.1, we extend the analysis in [54] to our maximum likelihood setting, and find that the estimation error from *N* samples is given by

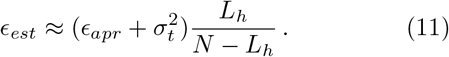

This expression is intuitively sensible: in the limit of infinite data, *N* → ∞, the estimation error vanishes, and in the opposite limit, *N* → *L*_*h*_, the estimation error blows up due to overfitting.

If 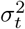 is not too small, *ϵ*_*est*_ is a monotonically increasing function of *L*_*h*_, as shown in Fig. 3 (green line). In particular, when *L*_*h*_ is significantly smaller than the number of training samples, *N, ϵ*_*est*_ is a linearly increasing function of *L*_*h*_, which is consistent with classical model selection theory [4, 12]. As *L*_*h*_ approaches *N*, the estimation error increases, and it goes to infinity when *L*_*h*_ = *N*, because the matrix on the right hand side of Eq. (7) becomes singular at that point.

Inserting *ϵ*_*est*_ from Eq. (11) into Eq. (4), the generalization error under MLE is

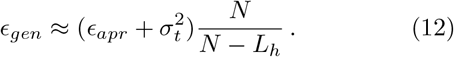

The generalization error typically has a nontrivial global minimum as a function of *L*_*h*_, as shown in Fig. 3 (red line). Moreover, the analytically estimated optimal hidden layer size, 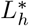, closely matches its estimation from numerical simulations (solid vs dotted vertical lines in Fig. 3).

### C. Optimal hidden layer size

By minimizing the generalization error, Eq. (12) with respect to *L*_*h*_ (with the approximation error given by Eq. (9)), we can find the optimal hidden layer size, 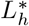, as a function of the input layer size, *L*_*x*_. As shown in Fig. 4A, 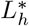 has three different scalings. That is because only one term at a time in Eq. (9) is sensitive to *L*_*h*_: the second term if *L*_*h*_ ∼ *𝒪* (1); the third term if *L*_*h*_ ∼ *𝒪* (*L*_*x*_) and the fourth term if 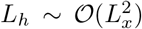. However, even considering one term at a time, minimizing Eq. (12) with respect to *L*_*h*_ is nontrivial, in large part because of the dependence on *N*. Details of the minimization are, therefore, left to SI, §5.1; here we simply summarize the results.

**FIG. 4.**
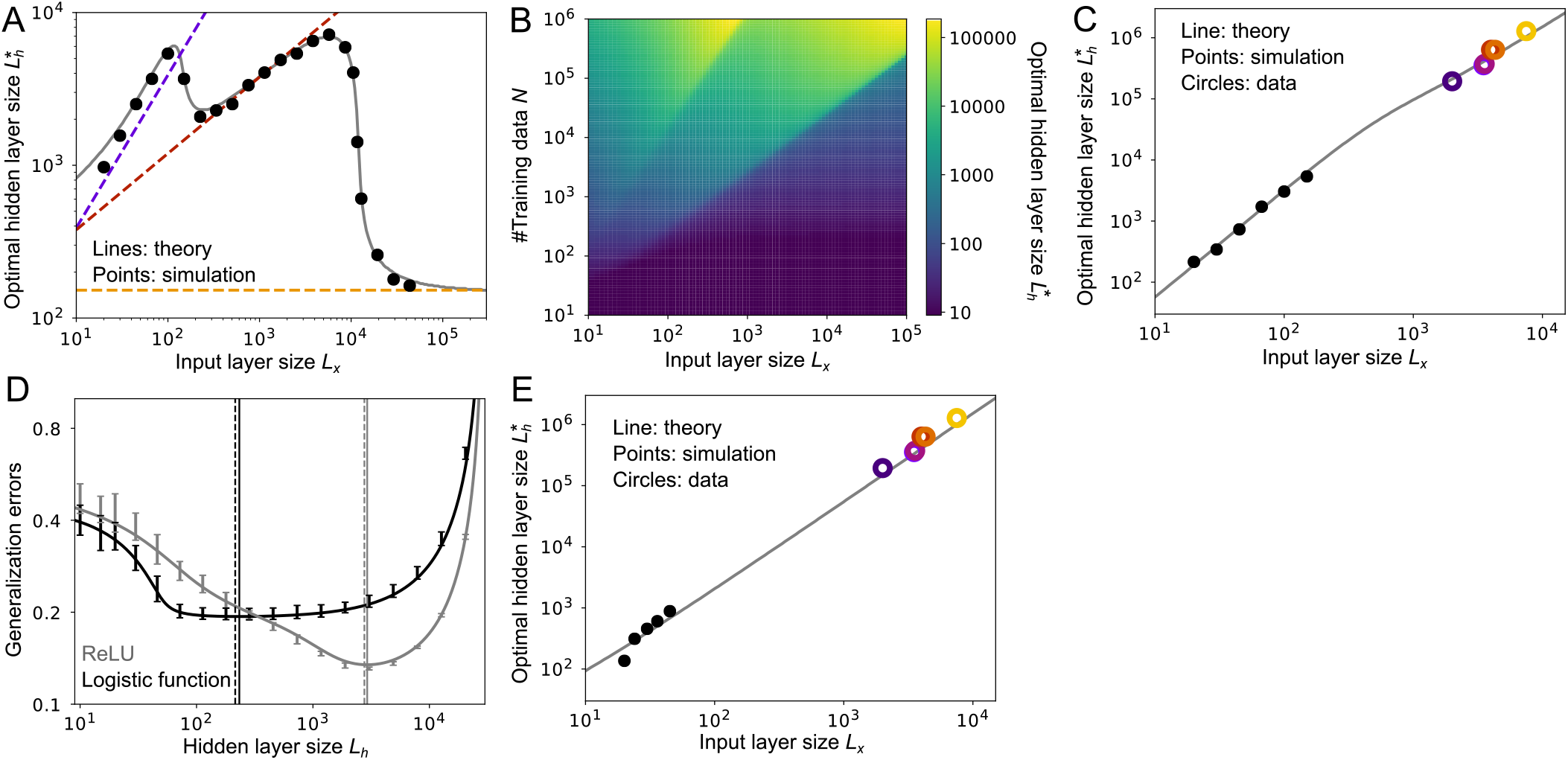
Model behavior under maximum likelihood estimation. **A)** Relationship between the input layer size, *L*_*x*_, and the optimal hidden layer size, 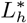, with a fixed sample size (*N* = 30000). Gray lines are found by optimizing Eq. (12) with respect to *L*_*h*_; dotted lines are the asymptotic expression derived in SI §5.1. **B)** Optimal hidden layer size, 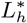, as a function of the input layer size, *L*_*x*_, and the sample size, *N*, from Eq. (12). **C)** Scaling with 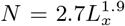. Gray line is theory; black points are from simulations; colored circles are the experimental data from Fig. 1A. Simulations were done only for low *L*_*x*_, due to the computational cost of the simulations when *L*_*x*_ is large. **D**) Relationship between the hidden layer size, *L*_*h*_, and the generalization error, *ϵ*_*gen*_, under the logistic activation function (black), and ReLU (gray), at *L*_*x*_ = 50 and *N* = 30000. Lines are theory; bars are from simulations. Error bars are the standard deviation over 10 simulations. **E**) Scaling for the logistic activation function with 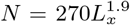. Gray line is theory; black points are from simulations; colored circles are the experimental data from Fig. 1A. As in panel C, simulations were done only for low *L*_*x*_, due to the computational cost of the simulations when *L*_*x*_ is large. As in Fig. 3, the teacher network had a hidden layer size of 500, used a ReLU nonlinearity, and the noise was set to 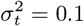.

The optimal hidden layer size, 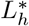, roughly follows one of the three dotted lines in Fig. 4A, depending on the value of *L*_*x*_. When the input layer size, *L*_*x*_, is small compared to *N*, 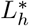 is linear in *L*_*x*_ (purple line in Fig. 4A); when *L*_*x*_ is comparable to *N, L*_*h*_ scales as the square root of *L*_*x*_ (red line); and when *L*_*x*_ is larger than *N, L*_*h*_ stays constant as *L*_*x*_ changes (orange line). This last scaling is reasonable because when the input layer is wide enough, expansion in the hidden layer is unnecessary. In all regions, 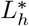 shows a square-root dependence on *N*, as suggested from previous studies [6, 8]. To further illustrate the dependence of 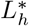 on *L*_*x*_ and *N*, in Fig. 4B we plotted the optimal hidden layer size versus these two quantities. This plot indeed shows three distinct phases separated by the lines *L*_*x*_ ∝ *N* and 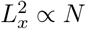.

Figure 4B shows that the scaling relationship between 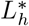 and *L*_*x*_ depends on *N*. Thus, to determine scaling across species, we need to know how *N* scales with *L*_*x*_ across species. We cannot directly measure *N*, which is the total amount of reward/teaching signal an animal experiences in its lifetime. However, we expect that *N* scales linearly with the duration of learning, so we can use that as a proxy for *N*. Among the six mammalian species, maximum longevity scales approximately as 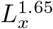 (SI, Fig. S1A; longevity data from AnAge database [55]). Alternatively, if we assume that learning happens mostly during the developmental period, here defined as the period from weaning to sexual maturation, a similar trend is observed, but with a slightly different exponent: duration from the time of weaning to sexual maturation scales approximately as 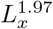 (SI, Fig. S1B).

Given these observations, we assumed 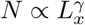 with *γ* between 1.5 and 2. When we did that, we found a clear scaling law between *L*_*x*_ and 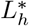 that spans more than three orders of magnitude. When we used 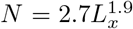, the model reproduced the 3/2 scaling observed in the mammalian olfactory system (Fig. 4C). (Other values of *γ* gave slightly different scaling; see SI, Fig. S2A.)

In the above examples, we used ReLU for both teacher (*g*_*t*_) and student (*g*_*s*_), but this matching (*g*_*t*_ = *g*_*s*_) is a rather strong assumption. To check the robustness of our results over the choice of the activation functions *g*, we used a logistic function *g*_*s*_ while keeping *g*_*t*_ as a ReLU. We found that the generalization error is minimized at a smaller hidden layer size compared to the ReLU student networks (black vs gray line in Fig. 4D; see SI §7.2 for details), primarily because large expansion is less helpful when the activation functions of the teacher and student networks are different. Nevertheless, assuming 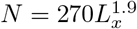, we obtain the experimentally observed 3/2 scaling law between 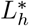 and *L*_*x*_ (Fig. 4E and Fig. S2B in SI).

### D. Stochastic gradient descent (SGD) learning

So far we have considered learning by maximum likelihood estimation (MLE). However, that is not the best choice when the hidden layer size, *L*_*h*_, is similar to the sample size *N*, as discussed above. In addition, batch learning is not particularly biologically plausible. Therefore, we consider online learning using stochastic gradient descent,

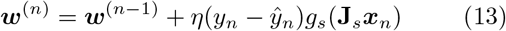

where ***w***^(*n*)^ is the readout weight after trial *n* and *η* is the learning rate. For online learning we consider minimization of the generalization error averaged over the lifetime of the organism, not the final error; that is because the fitness of an animal is much better characterized by the average proficiency during its lifetime than the proficiency at the end of its life.

Consistent with previous results [13], the learning rate that enables the fastest decay of the error is (see SI §4.2 for details)

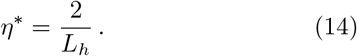

For this learning rate, the estimation error after *N* training samples is given approximately by (see SI §4.2, especially Eq. (91))

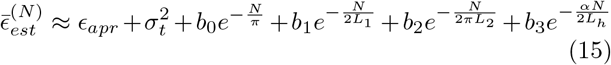

where

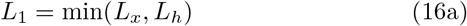

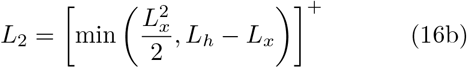

(recall that []^+^ is the rectified linear function). The coefficients *b*_0_, *b*_1_, *b*_2_ and *b*_3_ depend on *L*_*h*_, but not on *N*, and *α* (last term) is the same constant that appeared in Eq. (9).

The behavior of the estimation error under SGD is different than under MLE, Eq. (11), in two ways. First, for MLE, the estimation error goes to 0 as *N* → ∞; for SGD, it asymptotes to a constant. That is because we used a fixed learning for the SGD update rule rather than letting it decay, as would be necessary to reduce the estimation error to zero [56]. Second, for MLE the estimation error diverges as *L*_*h*_ approaches *N*, whereas for SGD it remains finite. That is because of the online nature of SGD, which precludes overfitting.

As expected from Eq. (15), the estimation error as a function of the number of training samples, *N*, exhibits three components, all decaying with different timescales (Fig. 5B). The timescales of these, *L*_1_, *L*_2_, and *L*_*h*_, are non-decreasing functions of *L*_*h*_, as shown in Fig. 5A. Thus, larger *L*_*h*_ means slower decay, as can be seen in Fig. 5B. Therefore, unless *L*_*h*_ is small (where the estimation error decreases because the coefficients *b*_*q*_ depend on *L*_*h*_), the lifetime average error increases with *L*_*h*_, as can be seen in the green line in Fig. 5C. Notably, though, because the estimation error remains finite as *L*_*h*_ → *N*, the lifetime average error does not diverge – in sharp contrast to maximum likelihood estimation, where it does diverge (compare the green line in Fig. 3 versus Fig. 5C). Because the approximation error decreases monotonically (blue line in Fig. 5C), the lifetime average generalization error (red line in Fig. 5C) tends to have a nontrivial global minimum.

**FIG. 5.**
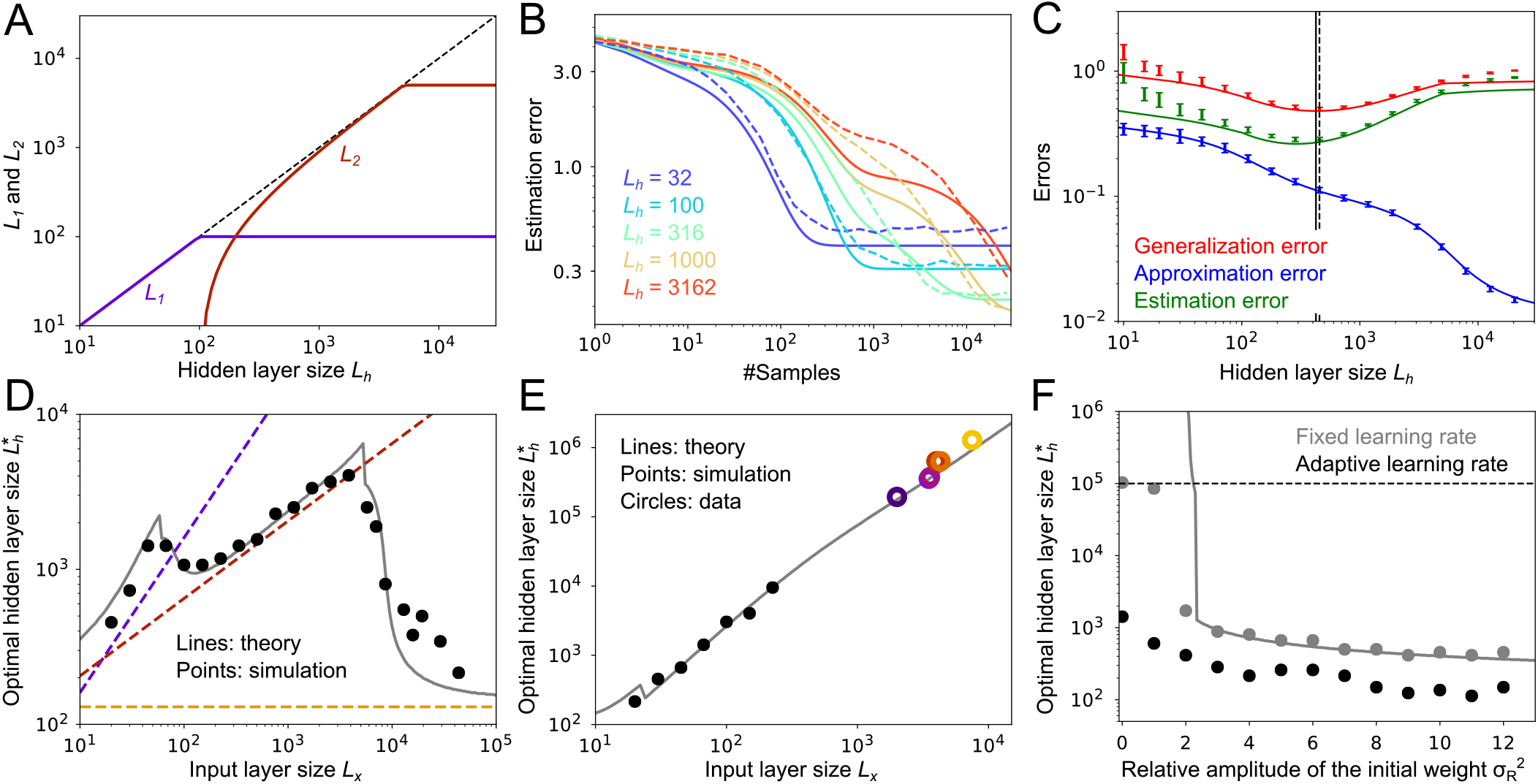
Model behavior under stochastic gradient descent. **A)** Hidden layer size dependence of the decay time constant *L*_1_ and *L*_2_, with *L*_*x*_ = 100. **B)** Dynamics of the estimation error under various hidden layer sizes, *L*_*h*_. Dashed lines: simulations; solid lines: theory. **C)** The cumulative generalization error, approximation error, and cumulative estimation error under various hidden layer sizes, *L*_*h*_, at *N* = 30000. **D)** Optimal hidden layer size, 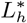, with *N* = 100000. Dotted lines are asymptotic scaling (see SI §5.2). **E)** Optimal hidden layer size, 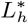, with 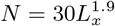. Gray line is theory; black points are from simulations; colored circles are the experimental data from Fig. 1A. As in Fig. 4, simulations were done only for low *L*_*x*_, due to the computational cost of the simulations when *L*_*x*_ is large. **F)** Optimal hidden layer size, 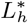, for various initial weight amplitudes, 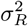, and *N* = 30000. Gray: fixed learning rate; black: adaptive learning rate. Lines are theory and dots are simulations. The initial readout weights were sampled from 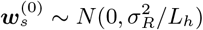. The horizontal dotted line represents the cutoff of 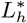 in the numerical simulations, meaning that at 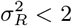, under a fixed learning rate 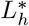 is larger than 10^5^. In panels A-C and F we set the input layer size to *L*_*x*_ = 100. As in Fig. 3, the teacher network had a hidden layer size of 500, used a ReLU nonlinearity, and the noise was set to 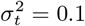.

As with MLE learning, under a fixed sample size *N* the optimal hidden layer size, 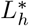, shows three different scalings (dotted lines in Fig. 5D). This is because the approximation error decreases with three distinct phases (Eq. (9)). As a result, we observe effectively the same structure in SGD that we saw in MLE (Fig. 5D vs Fig. 4A), although the theoretical prediction at large *L*_*x*_ under SGD does not match quite as well as under MLE. But by introducing the scaling 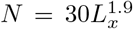, the experimentally observed scaling law in Fig. 1A is again reproduced (Fig. 5E).

In the model, we initialized the readout weights at relatively large values 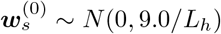,. If the weights are instead initialized to small values, the optimal hidden layer size 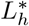 diverges to infinity (gray line and points in Fig. 5F). This is partially because the fixed learning rate (Eq. (14)), employed for analytical tractability, causes poor convergence at small *L*_*h*_. If an adaptive learning rate, *η*_*n*_ = 2*/* max(*L*_*h*_, *n*), is used instead [16, 57], the cumulative generalization error is optimal at a finite hidden layer size even when the initial readout weights are zero (black points in Fig. 5F). Although the optimal hidden layer size, 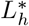, goes up as the initial weight amplitude 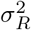 becomes smaller (Fig. 5F), the cumulative error becomes smaller under both fixed and adaptive learning rate (Fig. S3 in SI), due to smaller initial error.

### E. Evolutionary constraints

The results so far indicate that developmental constraints explain the scaling law observed in the mammalian olfactory system. However, our analysis also revealed that developmental constraints alone do not explain the 7/2 power law scaling observed in the invertebrate olfactory circuit, suggesting the presence of additional principles. The primary candidate is a constraint on the genetic budget an animal can use to specify the olfactory circuit. We refer to this as an evolutionary constraint. Because both the number of protein-encoding genes and the total size of the genome tends to be similar across species [58], we assume that the genetic budget for the specification of olfactory circuitry is similar among the insects listed on Fig. 1B.

Inspired by the insect olfactory circuitry, we consider a two-pathway model, in which projection neurons extend connections to both lateral horn neurons and Kenyon cells (Fig. 6A), and the output is

**FIG. 6.**
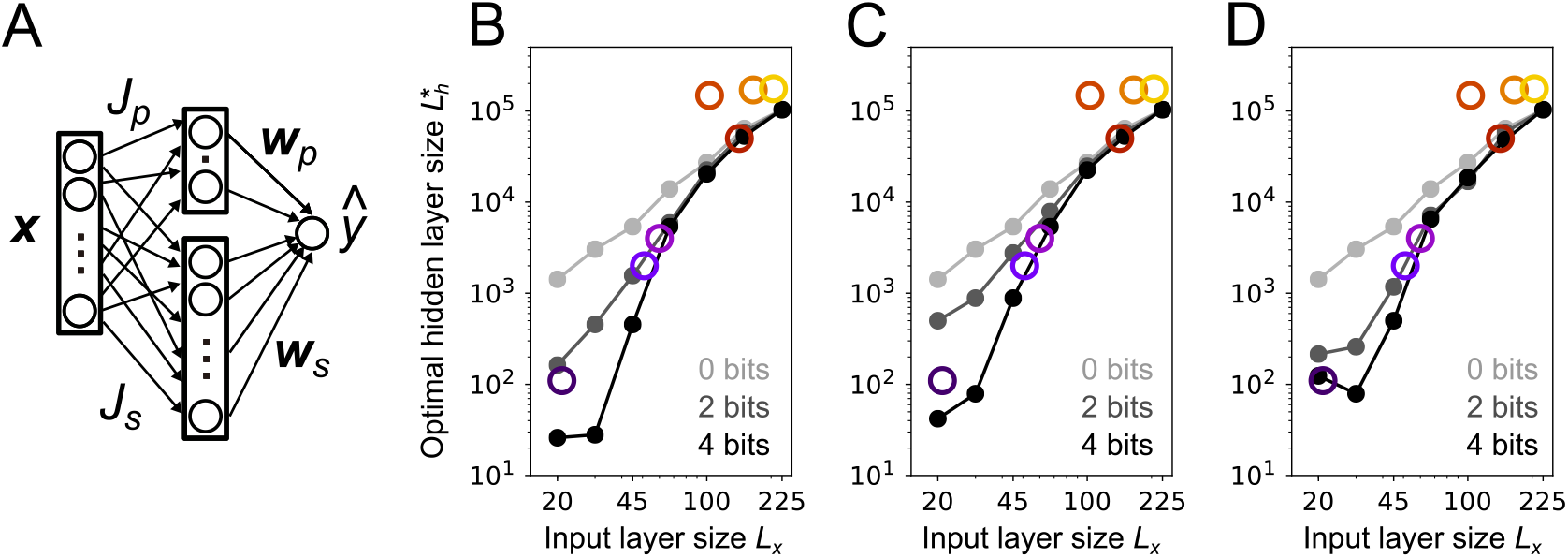
**A)** Schematic of the two pathway model. **B-D)** Optimal layer size of the randomly initialized pathway ***w***_*s*_ *g*(**J**_*s*_***x***) under different model settings. **B)** Low bit synapses were achieved by adding Gaussian noise to **J**_*p*_ and ***w***_*p*_. **C)** Low bit synapses were achieved by discretizing **J**_*p*_ and ***w***_*p*_. **D)** Low bit synapses were achieved by adding noise on **J**_*p*_ and ***w***_*p*_ as in B, but ***w***_*p*_ was additionally learned from training samples using SGD (see SI, Eq. (145)). In panels B-D, for the teacher network we used we had a hidden layer size of 500, used a ReLU nonlinearity, set the to 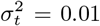, and used 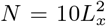 trials. For *s*_*b*_ = 2 bits we used *G* = 2000, while for *s*_*b*_ = 4 bits we used *G* = 4000, and for *s*_*b*_ = 0 bit, we simply removed the hard-wired pathway. The width of the hardwired intermediate layer, *L*_*p*_, was found from Eq. (18): *L*_*p*_ = *G/s*_*b*_(*L*_*x*_ + 1), rounded up to an integer. See SI §6 and §7.4 for details.

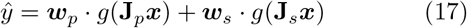

where ***w***_*p*_ *g*(**J**_*p*_***x***) is the pathway through lateral horn neurons. Although lateral horn neurons do not directly project to mushroom body output neurons, the two pathways eventually converge in the pre-motor area [26], where the output *ŷ* might be represented. Because connections between projection neurons and lateral horn neurons tend to be stereotypical [20, 47], we assumed they were optimized on evolutionary timescales. We thus tuned the values of **J**_*p*_ and ***w***_*p*_ while constraining the total information required for specifying the weights (see SI §6). In contrast, **J**_*s*_ was initialized randomly and fixed, and ***w***_*s*_ was learned with adaptive SGD. Note that the initialization of **J**_*s*_ and ***w***_*s*_ should require very little genetic information, compared to the hard-wired projection neuron-to-lateral horn neuron pathway. Using *L*_*p*_ to denote the number of lateral horn neurons, under a genetic information budget *G*, the amount of information encoded in the initial condition of **J**_*p*_ and ***w***_*p*_ is bounded by

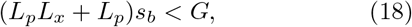

where *s*_*b*_ is the number of bits per synapse. The first term is the number of bits needed to specify **J**_*p*_; the second is the number needed to specify ***w***_*p*_. When only the presence/absence of connections is determined genetically, *s*_*b*_ is at most 1 bit; additional bits are needed if the weights are specified as well. Under a fixed budget, *G*, the number of bits per synapse, *s*_*b*_, is bounded by *G/L*_*x*_*L*_*p*_, suggesting that as the input layer size, *L*_*x*_, increases, tuning of **J**_*p*_ and ***w***_*p*_ have to be more coarsegrained. In particular, in the mammalian olfactory system where *L*_*x*_ ∼ 10^3^, the hard-wired pathway should play a minor role unless *G >* 10^4^. Indeed, except for encoding of pheromone signals, evidence of hard-wired connections in the mammalian olfactory circuits is limited [59]. For invertebrates, which have far fewer glomeruli, hard-wired pathways should be far more important. As the effect of the genetic budget, *G*, is difficult to characterize analytically, we numerically investigate its effect.

When we allowed information about the weights to be transmitted genetically, subject to the constraint given in Eq. (18), we found that the optimal Kenyon cell population size, *L*_*h*_, was much smaller than the circuit without the projection neuron-to-lateral horn neuron pathway (compare 0 bit lines to 2 and 4 bit lines in Fig. 6B-D), leading to steeper scaling. Note that the two estimates became close at large *L*_*x*_ (as predicted above). Thus, for mammals, which have *L*_*x*_ ∼ 10^3^, genetic information has a negligible effect on scaling. In particular, we found that by setting *s*_*b*_ = 2, and *G* = 2000, the 7/2 scaling observed among insects is approximately reproduced (dark gray line in Fig. 6B). The predicted curve saturates at square scaling around *L*_*x*_ ≈ 150, resulting in under-estimation of the Kenyon cell population in bees and cockroaches. This trend was observed under a different implementation of low-bit synapses (Fig. 6C), and even when ***w***_*p*_ was additionally trained with SGD from finely-tuned initial weights (Fig. 6D).

## III. DISCUSSION

In this work, we modeled the olfactory circuit of both mammals and insects as a three layer feedforward network, and asked how the number of neurons in the hidden layer scales with the number neurons in the glomerular (i.e., input) layer. We hypothesized that the scarcity of labeled signals (e.g., reward and punishment) provides a crucial constraint on the hidden layer size. We showed analytically, and confirmed with simulations, that this hypothesis robustly explains the experimentally observed 3/2 scaling found in the mammalian olfactory circuit, both under maximum likelihood (Fig. 4) and stochastic gradient descent (Fig. 5) learning. (Here “3/2 scaling” means the number of neurons in the hidden layer is proportional to the number of glomeruli to the 3/2 power.) This hypothesis alone does not, however, explain the 7/2 scaling found in the olfactory circuit of insects. But by considering the fact that genetic information used for constructing hard-wired olfactory connections is limited, we recovered the 7/2 scaling law (Fig. 6), without disrupting the 3/2 scaling law in mammals.

The 3/2 power in the scaling law we derived for mammals comes from two factors. First, when the number of training samples is fixed, the optimal population size of the piriform cortex increases as the number of glomeruli increases, unless the number of glomeruli is very large (Figs. 4A and 5D). Second, the optimal population size of the piriform cortex also increases with the number of training samples (Fig. 4B). Because species with more glomeruli tend to live longer and experience more samples (Fig. S1), this sample size dependence causes an additional scaling between the number of glomeruli and the piriform population size. From these two factors, the optimal intermediate layer size scales supra-linearly on number of the glomeruli (Figs. 4C, E, and 5E). Because of the dependence on the number of training samples, *N*, the power in the scaling law is not fixed at 3/2. In fact, depending on how *N* scales with the input layer size, *L*_*x*_, theoretically any scaling is possible (SI §5). The 3/2 scaling we found was because in mammals, lifetime scales approximately quadratically with the number of glomeruli (SI §1.2, Table 2).

The three layer feedforward neural network with random fixed hidden weights is a class of neural networks that is widely studied from both biological [3, 37, 38, 60] and engineering [61, 62] perspectives. Under batch learning, the upper bound on the approximation error for this network structure is known for a large class of the target functions [63, 64], yet these bounds are often too loose in practice. Here, we instead derived the average approximation error (SI §3). This allowed us to derive, analytically, accurate estimates of the optimal hidden layer size. The behavior of the estimation error is also well characterized in the large sample size limit (*N* → ∞ while *L*_*x*_, *L*_*h*_ *<* ∞) [12, 65], but this limit is not a good approximation of an over-parameterized neural network. On the other hand, the characteristics of the error at the large parameter space limit (number of synapses ∝ *N* as *N*→ ∞) remains mostly elusive, except for linear regression [54] (see also §4.1). Similarly, model selection in neural networks was previously studied mostly in the large sample size limit [7, 66]. The upper bound on the network size was also studied from VC (Vapnik-Chervonenkis) theory [5], and the minimum description length principle [6].

Learning dynamics in neural networks under SGD has also been widely studied [13, 49, 57]. In particular, recent results suggest that over-parameterization of a neural network does not harm the generalization error under both full-batch and stochastic gradient descent learning [67–70]. Here, though, we focused on the cumulative error, not the error at the end of training, as the former is more relevant to the fitness of the species. Under this objective function, over-parameterization does tend to harm performance, because learning becomes slower (Fig. 5B), even under an adaptive learning rate (Fig. 5F). Nevertheless, we found that if the initial weights are set to very small values and the learning rate is fixed, having infinitely many neurons in the hidden neuron minimizes the cumulative error (Fig. 5F), suggesting that over-parameterization is not always harmful, even when the cumulative error is the relevant cost function.

Scaling laws are ubiquitous in the brain. For instance, the number of neurons in the primary visual cortex scales with the 3/2 power against the population size of the LGN [71], while the number of neurons in the cerebral cortex is linear in the total number of neurons in the cerebellum [72]. Given the anatomical similarity between the olfactory circuit and cerebellum [3], our methodology should be directly applicable to understanding the latter scaling. But it is not limited to olfactory-like structures; it could be applied, possibly with some modifications, anywhere in the brain, and has the potential to provide insight into circuit structure in general.

## Code availability

The source codes of the simulations and the data analysis are deposited at Github (https://github.com/nhiratani/olfactorydesign).

## Acknowledgement

This work was supported by the Gatsby Charitable Foundation and the Wellcome Trust (110114/Z/15/Z).

## Competing interests

The authors declare no competing interests.

## Notes

### Competing Interest Statement

The authors have declared no competing interest.

