## Supplementary information for "Developmental and evolutionary constraints on olfactory circuit selection"

#### Supplementary figures

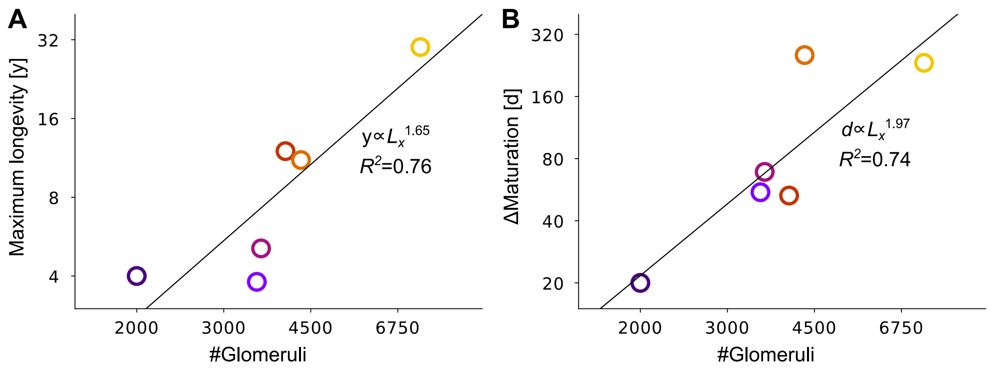


**Figure S1:** Scaling of duration of learning for vertebrates with the number of glomeruli. **A)** Maximum longevity versus number of glomeruli. **B)** Average duration from weaning to sexual maturation versus number of glomeruli. Color code is the same as in Fig. 1A.

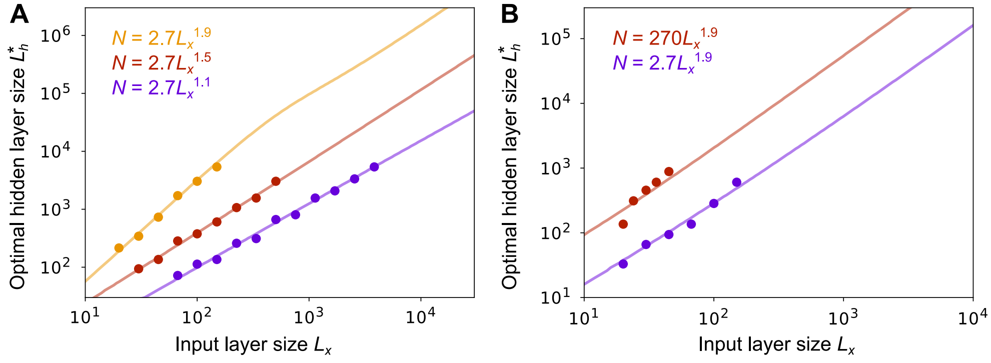


**Figure S2:** Scaling between the optimal hidden layer size and the input layer size under maximum likelihood estimation. **A)** Scaling under various exponents relating $N$ to $L_x$. Here, the activation function of both student and teacher networks are ReLU. **B)** Scaling when the student activation function is the logistic function while the teacher function is ReLU. In both panels, points are simulations and lines are analytical results.

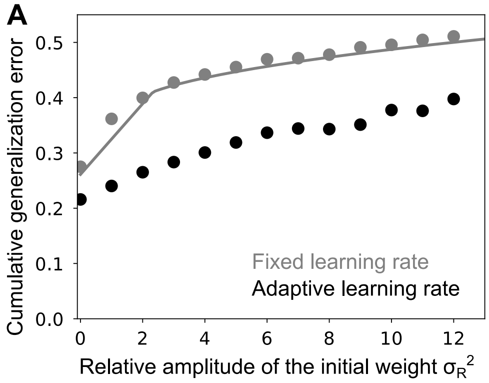


**Figure S3:** The cumulative generalization error $\epsilon_{cg}^N$ under various initial weight amplitude $\sigma_R^2$, when learning is performed with a fixed learning rate (gray), and an adaptive learning rate (black). The hidden layer size, $L_h$, was set to the optimal value estimated numerically in Fig. 5F. The initial projection weights were sampled from $\boldsymbol{w}_s^{(0)} \sim N(0, \sigma_R^2/L_h)$.

### 1 Data analysis

#### 1.1 Scaling in the invertebrate olfactory circuitry

Table 1 gives the number of glomeruli and Kenyon cells that were used to make Fig. 1B, along with the sources. All numbers are estimates for one hemisphere. It should be noted that the data is not well controlled. For instance, in some cases only the total number of mushroom body neurons are available, instead of the number of Kenyon cells, and different experimental techniques were used for different animals. Moreover, for locusts we used the number of olfactory receptor genes as an estimate of the effective number of glomeruli, because they have a unique micro-glomeruli structure which makes a direct comparison difficult [1]. There is usually a one-to-one correspondence between the number of olfactory receptor genes and the number of glomeruli in the invertebrate olfactory system, so this should be a good proxy of the effective number of glomeruli [2, 3]. For the total number of glomeruli, a comprehensive review is available [4]. Further explanation of the data for each species is given below.

| Species | #Glomeruli | #KC | ref(G) | ref(KC) |
|---|---|---|---|---|
| Drosophila Melanogaster (larva) | 21 | 110 | [5, 6] | [7] |
| Drosophila Melanogaster (adult) | 51 | 2000 | [8] | [9] |
| Moth (Spodoptera litoralis) | 60 | 4000 | [10] | [11] |
| Locust | 142* | 50000 | [12] | [13] |
| Apis mellifica (drone) | 103 | 148000 | [14] | [15] |
| Apis mellifica (worker) | 165 | 170000 | [14] | [15] |
| Cockroach (Periplaneta americana) | 205 | 175000 | [16] | [17] |

Table 1. Number of glomeruli and Kenyon cells (KCs) for seven invertebrate species. *For locusts, the number of olfactory receptor genes is shown.

**Drosophila Melanogaster (larvae)**

Recent studies suggest that fruit fly larvae have a well functioning olfactory system, though the corresponding circuit is much smaller than that of adults [18]: there are only 21 glomeruli [5, 6], and about 110 Kenyon cells [7].

**Drosophila Melanogaster (adults)**

The adult fruit fly is the best studied species of insect. Their olfactory system contains 51 glomeruli that project to the antennal lobe [8], and around 2200 mushroom body neurons, of which about 2000 are Kenyon cells [9].

**Moth (Spodoptera litoralis)**

Several species of moth have been studied, and they all have around sixty glomeruli [4], including Spodoptera litoralis [10]. Less is known about the number of Kenyon cells, but one study found that the mushroom body of Spodoptera litoralis contains around 4000 of them [11].

**Locust**

Unlike most invertebrates, the locust olfactory circuit has a micro-glomeruli structure [1], meaning each micro-glomerulus receives input from multiple types of olfactory receptor neurons. This makes it difficult to compare with other species. However, we can use the number of olfactory receptor genes as a proxy, as discussed above. Under this assumption, the number is about 142 [12]. The number of Kenyon cells is estimated to be around 50000 [13].

**Apis mellifica (drone)**

Due to the caste system of honey bees, male honey bees (drones) do not engage in foraging or colony protection. Correspondingly, they have a smaller number of glomeruli compared to worker bees (103 vs 165 [14]), despite their larger body size. Their mushroom body is estimated to contain about 148000 neurons [15]. As shown in Figure 1B, the drone bee is a clear outlier from the scaling law. However, this is understandable considering its unique ecological niche.

**Apis mellifica (worker)**

As mentioned above, worker bees have around 165 glomeruli [14], and the number of neurons in the mushroom body is around 170000 [15]. Note that the mushroom body of the worker honey bee is known to take part in visual navigation as well as olfaction [19].

**Cockroach (Periplaneta americana)**

Cockroaches are known to have excellent olfactory discrimination and learning ability [20]. They have about 205 glomeruli [16] and around 175000 Kenyon cells [17].

##### 1.2 Correlation between the number of glomeruli and the duration of learning

The maximum longevity and average time from weaning to sexual maturation can be estimated from the AnAge database [21]. These are summarized in Table 2, for the six mammalian species used in our analysis (Fig. 1A) [22]. For species with different sexual maturation times for males and females, we took the average.

| Species | common name | longevity (y) | weaning (d) | sexual maturity (d) | $\Delta$maturation (d) |
|---|---|---|---|---|---|
| Mus musculus | mice | 4.0 | 22 | 42 | 20 |
| Rattus norvegicus | rats | 3.8 | 25 | 80 | 55 |
| Monodelphis domestica | opossums | 5.1 | 53 | 122 | 69 |
| Cavia porcellus | guinea pig | 12 | 18 | 71 | 53 |
| Mustela putorius | ferrets | 11.1 | 63 | 317 | 254 |
| Felis catus | cats | 30 | 56 | 289 | 233 |

Table 2. Maximum longevity, the time of weaning and of sexual maturation, and the difference of the latter two, denoted $\Delta$maturation, for the six mammalian species shown in Fig. 1A. y = year; d = day.

Supplementary Figs. S1A and S1B were obtained by plotting the data above (Table 2) against the number of glomeruli in Table 3 of [22].

#### 2 Model setting

We consider a three layer student-teacher model. We assume that the generative model of the environment (teacher model) is

$$y = \boldsymbol{w}_t \cdot g_t(\boldsymbol{J}_t \boldsymbol{x}) + \sigma_t \xi, \tag{1}$$

where $\boldsymbol{x} \in \Re^{L_x}$ is the olfactory input as, $y \in \Re$ is the associated reward/valence/label, $\boldsymbol{w}_t \in \Re^{L_t}$ and $\boldsymbol{J}_t \in \Re^{L_t \times L_x}$ are random matrices with elements drawn from a zero mean Gaussian,

$$w_j^t \sim \mathcal{N}(0, 1/L_t) \tag{2a}$$

$$J_{ij}^t \sim \mathcal{N}(0, 1/L_x), \tag{2b}$$

and $\sigma_t \xi$ is the teacher noise, which reflects the probabilistic correspondence between $\boldsymbol{x}$ and $y$. Here $\xi$ is a zero mean, unit variance Gaussian random variable. Throughout the text we use bold capital letters to denote matrices and bold small letter for vectors. As in the main text, vectors are defined as column vectors, and a superscript $T$ denotes transpose (indicating a row vector). For readibility, we use a dot product to denote the inner product between two vectors.

The olfactory circuit (student model) needs to mimic the teacher model to predict the reward/valence/label, $y$, given olfactory input, $\boldsymbol{x}$. We approximate this circuit by a three-layer feedforward network,

$$\hat{y} = \boldsymbol{w}_s \cdot g_s(\boldsymbol{J}_s \boldsymbol{x}), \tag{3}$$

where $\boldsymbol{w}_s \in \Re^{L_h}$ and $\boldsymbol{J}_s \in \Re^{L_h \times L_x}$. We assume that $\boldsymbol{J}_s$ is fixed and random with, as for the teacher network, entries drawn from a zero mean Gaussian,

$$J_{ij}^s \sim \mathcal{N}(0, 1/L_x)\,. \tag{4}$$

The goal of learning is to tune the projection weights, $\boldsymbol{w}_s$, based on the samples generated from the teacher model, $D_N = \{\boldsymbol{x}_n, y_n\}_{n=1}^N$. For analytical tractability we assume that the olfactory inputs, $\boldsymbol{x}$, are sampled from an uncorrelated Gaussian distribution, $\boldsymbol{x} \sim \mathcal{N}(0, \boldsymbol{I})$.

For the objective function, we use the mean squared error averaged over the input distribution, $p(\boldsymbol{x})$, and the teacher noise distribution, $p(\xi)$,

$$\epsilon_{gen} \equiv \left\langle \left[\boldsymbol{w}_t \cdot g_t(\boldsymbol{J}_t \boldsymbol{x}) + \sigma_t \xi - \boldsymbol{w}_s \cdot g_s(\boldsymbol{J}_s \boldsymbol{x})\right]^2 \right\rangle_{p(\boldsymbol{x},\xi)}. \tag{5}$$

Under this loss function, the optimal projection weight, $\boldsymbol{w}_s^*$, is given by the standard expression for linear regression,

$$\boldsymbol{w}_s^* \equiv \left\langle g_s(\boldsymbol{J}_s \boldsymbol{x}) g_s(\boldsymbol{J}_s \boldsymbol{x})^T \right\rangle^{-1} \left\langle g_s(\boldsymbol{J}_s \boldsymbol{x}) g_t(\boldsymbol{J}_t \boldsymbol{x})^T \right\rangle \boldsymbol{w}_t. \tag{6}$$

When learning is unbiased, the generalization error divides cleanly into an approximation error and an estimation error,

$$\epsilon_{gen} = \sigma_t^2 + \left\langle \left( [\boldsymbol{w}_t \cdot g_t(\boldsymbol{J}_t\boldsymbol{x}) - \boldsymbol{w}_s^* \cdot g_s(\boldsymbol{J}_s\boldsymbol{x})] + [(\boldsymbol{w}_s^* - \boldsymbol{w}_s) \cdot g_s(\boldsymbol{J}_s\boldsymbol{x})] \right)^2 \right\rangle_{p(\boldsymbol{x},\xi)} = \sigma_t^2 + \epsilon_{apr} + \epsilon_{est}, \tag{7}$$

where

$$\epsilon_{apr} \equiv \left\langle [\boldsymbol{w}_t \cdot g_t(\boldsymbol{J}_t\boldsymbol{x}) - \boldsymbol{w}_s^* \cdot g_s(\boldsymbol{J}_s\boldsymbol{x})]^2 \right\rangle_{p(\boldsymbol{x})} \tag{8a}$$

$$\epsilon_{est} \equiv \left\langle [(\boldsymbol{w}_s - \boldsymbol{w}_s^*) \cdot g_s(\boldsymbol{J}_s\boldsymbol{x})]^2 \right\rangle_{p(\boldsymbol{x})}. \tag{8b}$$

The approximation error, $\epsilon_{apr}$, depends only on the architecture, while the estimation error, $\epsilon_{est}$, depends on both the choice of the learning method and the number of trials, $N$. Below we derive approximate analytical expressions for both $\epsilon_{apr}$ and $\epsilon_{est}$, starting with the former.

### 3 Approximation error

To reduce clutter, we make the definitions

$$\boldsymbol{g}_t \equiv g_t(\boldsymbol{J}_t\boldsymbol{x}) \tag{9a}$$

$$\boldsymbol{g}_s \equiv g_s(\boldsymbol{J}_s\boldsymbol{x}). \tag{9b}$$

In terms of these quantities, the approximation error is

$$\epsilon_{apr} = \boldsymbol{w}_t^T \left( \langle \boldsymbol{g}_t \boldsymbol{g}_t^T \rangle - \langle \boldsymbol{g}_t \boldsymbol{g}_s^T \rangle \boldsymbol{G}_s^{-1} \langle \boldsymbol{g}_s \boldsymbol{g}_t^T \rangle \right) \boldsymbol{w}_t \tag{10}$$

where the angle brackets represent an average over $p(\boldsymbol{x})$, the distribution of the input, and $\boldsymbol{G}_s$ is the uncentered hidden layer covariance matrix,

$$\boldsymbol{G}_s \equiv \langle \boldsymbol{g}_s \boldsymbol{g}_s^T \rangle. \tag{11}$$

Computing $\epsilon_{apr}$ is hard because it involves the inverse of the covariance matrix, $\boldsymbol{G}_s$. However, for the model we consider, the off-diagonal elements can be expanded in powers of $1/L_x^{1/2}$ (as we show below). We make use of this expansion to compute (approximately) the eigenvalues and eigenvectors of $\boldsymbol{G}_s$, and use those to find the inverse. That calculation is described next. In the bulk of the analysis we consider arbitrary nonlinear functions $g_s(\cdot)$ and $g_t(\cdot)$. In our numerical analysis we use ReLU and logistic functions.

##### Hidden layer covariance

The hidden layer covariance is computed by averaging over $\boldsymbol{x}$. Note, though, that wherever $\boldsymbol{x}$ appears it is multiplied by $\boldsymbol{J}_s$, so instead of averaging over $\boldsymbol{x}$ we can average over $\boldsymbol{u} \equiv \boldsymbol{J}_s\boldsymbol{x}$. Because $\boldsymbol{x}$ is Gaussian and white, $\boldsymbol{u}$ is also Gaussian, but it is correlated,

$$\boldsymbol{u} \sim N(0, \boldsymbol{J}_s\boldsymbol{J}_s^T). \tag{12}$$

We thus have

$$(\boldsymbol{G}_s)_{ij} = \int du_i du_j \, p(u_i, u_j) g_s(u_i) g_s(u_j) \tag{13}$$

where $p(u_i, u_j)$ is a correlated Gaussian distribution with variance $\sigma_i^2$ and correlation coefficient $\rho_{ij}$; these quantities are give by

$$\sigma_i^2 = (\boldsymbol{J}_s\boldsymbol{J}_s^T)_{ii} \approx \left\langle (\boldsymbol{J}_s\boldsymbol{J}_s^T)_{ii} \right\rangle_{p(\boldsymbol{J}_s)} = 1. \tag{14a}$$

$$\rho_{ij} = \frac{(\boldsymbol{J}_s\boldsymbol{J}_s^T)_{ij}}{\sigma_i \sigma_j} \approx (\boldsymbol{J}_s\boldsymbol{J}_s^T)_{ij}, \; i \neq j. \tag{14b}$$

We will use $\sigma_i^2 = 1$ in what follows.

Let us first consider the diagonal terms which, under the approximation that $\sigma_i^2 = 1$, are all the same,

$$\langle g_s(u_i)^2 \rangle \approx \int_{-\infty}^{\infty} \frac{du_i}{\sqrt{2\pi}} \exp\left(-\frac{u_i^2}{2}\right) g_s(u_i)^2 \equiv D_0^s. \tag{15}$$

For instance, if $g_s(\cdot)$ is ReLU, $D_0^s = 1/2$. Even when $g_s(\cdot)$ is a complicated function, $D_0^s$ can be evaluated numerically (see §7.2). For the off-diagonal terms ($i \neq j$), again under the approximation $\sigma_i^2 = 1$, we have

$$\langle g_s(u_i) g_s(u_j) \rangle \approx \int_{-\infty}^{\infty} \int_{-\infty}^{\infty} \frac{du_i du_j}{2\pi (1-\rho_{ij}^2)^{1/2}} \exp\left( -\frac{u_i^2 + u_j^2 - 2\rho_{ij} u_i u_j}{2(1-\rho_{ij}^2)} \right) g_s(u_i) g_s(u_j) \,. \tag{16}$$

Expansion of the $\rho$-dependent terms in Eq. (16) around $\rho_{ij} = 0$ gives

$$\frac{1}{\sqrt{1-\rho_{ij}^2}} \exp\left( -\frac{u_i^2 + u_j^2 - 2\rho_{ij} u_i u_j}{2(1-\rho_{ij}^2)} \right) = \exp\left( -\frac{u_i^2}{2} - \frac{u_j^2}{2} \right) \left[ 1 + u_i u_j \rho_{ij} + \frac{(1-u_i^2)(1-u_j^2)}{2} \rho_{ij}^2 + O(\rho_{ij}^3) \right]. \tag{17}$$

Consequently, when $i \neq j$, $\langle g_s(u_i) g_s(u_j) \rangle$ can be approximated as

$$\langle g_s(u_i) g_s(u_j) \rangle \approx C_0^{ss} + C_1^{ss} \rho_{ij} + C_2^{ss} \rho_{ij}^2, \tag{18}$$

where

$$C_0^{ss} \equiv \langle g_s(u) \rangle_{\mathcal{N}}^2 \tag{19a}$$

$$C_1^{ss} \equiv \langle u g_s(u) \rangle_{\mathcal{N}}^2 \tag{19b}$$

$$C_2^{ss} \equiv \frac{1}{2} \left\langle [1-u^2] g_s(u) \right\rangle_{\mathcal{N}}^2 \,. \tag{19c}$$

The subscript $\mathcal{N}$ indicates an average over a standard Normal: for any function $g(u)$,

$$\langle g(u) \rangle_{\mathcal{N}} \equiv \int_{-\infty}^{\infty} \frac{du\, e^{-u^2/2}}{(2\pi)^{1/2}} g(u) \,. \tag{20}$$

To estimate the size of $\rho_{ij}$, we note that its mean and variance are given by

$$\langle \rho_{ij} \rangle_{p(\boldsymbol{J}_s)} = \left\langle [\boldsymbol{J}_s \boldsymbol{J}_s^T]_{ij} \right\rangle_{p(\boldsymbol{J}_s)} = 0, \tag{21a}$$

$$\langle \rho_{ij}^2 \rangle_{p(\boldsymbol{J}_s)} = \left\langle \left( [\boldsymbol{J}_s \boldsymbol{J}_s^T]_{ij} \right)^2 \right\rangle_{p(\boldsymbol{J}_s)} = \frac{1}{L_x}. \tag{21b}$$

Consequently, the correlation, $\rho_{ij}$, is in the order of $1/L_x^{1/2}$. However, there are $L_h$ times more off-diagonal terms than diagonal terms in the matrix $\boldsymbol{G}_s$, so we cannot ignore the $\rho_{ij}$-dependent terms in Eq. (18) unless $L_h^2 \ll L_x$. Nevertheless, for $L_x \gg 1$ (the relevant limit in our analysis), the correlation satisfies $|\rho_{ij}| \ll 1$, suggesting that for large $L_x$, a second-order Taylor expansion in $\rho_{ij}$ should provide a good approximation to $\langle g(u_i) g(u_j) \rangle$. That's the approach we take here.

Combining the diagonal (Eq. (15)) and off-diagonal (Eq. (18)) terms, we can write the full covariance matrix as

$$\langle g_s(u_i) g_s(u_j) \rangle \approx D_0^s \delta_{ij} + (C_0^{ss} + C_1^{ss} \rho_{ij} + C_2^{ss} \rho_{ij}^2)(1 - \delta_{ij}) \,. \tag{22}$$

It is convenient to collect the diagonal terms, and to make the second-order term zero-mean, yielding

$$\langle g_s(u_i) g_s(u_j) \rangle \approx \delta_s \delta_{ij} + (C_0^{ss} + C_2^{ss} \langle \rho^2 \rangle) + C_1^{ss} \rho_{ij} + C_2^{ss} (\rho_{ij}^2 - \langle \rho^2 \rangle) \tag{23}$$

where $\langle \rho^2 \rangle$ is defined in Eq. (21b) and

$$\delta_s \equiv D_0^s - (C_0^{ss} + C_1^{ss} + C_2^{ss}) \,. \tag{24}$$

Strictly speaking, $\rho_{ij}$ is not defined at $i = j$, but we can choose it arbitrarily without changing the covariance matrix. We thus extend Eq. (14b) to include $i = j$, and write (again using $\sigma_i = 1$)

$$\rho_{ij} = (\boldsymbol{J}_s \boldsymbol{J}_s^T)_{ij} \,, \tag{25}$$

now valid for $i = j$ as well as $i \neq j$. Using this convention, it is straightforward to show that

$$\rho_{ij}^2 - \langle \rho^2 \rangle = \sum_{m=1}^{L_x} \sum_{l=m+1}^{L_x} M_{i,[m,l]}^s M_{j,[m,l]}^s \tag{26}$$

where $[m,l]$ is a compositional index, and $\boldsymbol{M}_s$ is an $L_h \times L_x(L_x-1)/2$ matrix,

$$M^s_{i,[m,l]} \equiv \sqrt{2} J^s_{im} J^s_{il} \,. \tag{27}$$

Combining these expressions, and using the fact that $\langle \rho^2 \rangle = 1/L_x$ (Eq. (21b)), which is small compared to 1, the covariance matrix simplifies to

$$\boldsymbol{G}_s \approx \delta_s \boldsymbol{I} + C_0^{ss} \boldsymbol{1}_h \boldsymbol{1}_h^T + C_1^{ss} \boldsymbol{J}_s \boldsymbol{J}_s^T + C_2^{ss} \boldsymbol{M}_s \boldsymbol{M}_s^T \tag{28}$$

where $\boldsymbol{1}_h \in \Re^{L_h}$ is a vector in which all the elements are one. The first two matrices in this expression are a scaled identity matrix and a matrix with the same value everywhere. The third, $\boldsymbol{J}_s \boldsymbol{J}_s^T$, is a Wishart matrix, so its eigenspectrum follows a Marchenko-Pastur distribution [23]; from Eq. (4) we see that the parameters of that distribution are $(\sigma^2 = 1, \lambda = L_h/L_x)$. Similarly, although the columns of $\boldsymbol{M}_s$ are not independent, given that they have zero correlation we assume that the eigenspectrum of $\boldsymbol{M}_s \boldsymbol{M}_s^T$ also follows a Marchenko-Pastur distribution; from Eq. (27) we see that the parameters are $(\sigma^2 = 1, \lambda = 2L_h/L_x^2)$.

Essentially identical analysis gives us the covariance between the hidden units of the teacher and student networks,

$$\langle \boldsymbol{g}_t \boldsymbol{g}_s^T \rangle \approx C_0^{ts} \boldsymbol{1}_t \boldsymbol{1}_h^T + C_1^{ts} \boldsymbol{J}_t \boldsymbol{J}_s^T + C_2^{ts} \boldsymbol{M}_t \boldsymbol{M}_s^T \,, \tag{29}$$

where $\boldsymbol{M}_t$ is an $L_t \times L_x(L_x-1)/2$ matrix analogous to $\boldsymbol{M}_s$,

$$M^t_{i,[k,l]} \equiv \sqrt{2} J^t_{ik} J^s_{il}, \tag{30}$$

and $C_0^{ts}, C_1^{ts}, C_2^{ts}$ are natural extensions of $C_0^{ss}, C_1^{ss}, C_2^{ss}$ (Eq. (19)), the difference being that one of the student averages becomes a teacher average,

$$C_0^{ts} \equiv \langle g_t(u) \rangle_{\mathcal{N}} \langle g_s(u) \rangle_{\mathcal{N}} \tag{31a}$$

$$C_1^{ts} \equiv \langle u g_t(u) \rangle_{\mathcal{N}} \langle u g_s(u) \rangle_{\mathcal{N}} \tag{31b}$$

$$C_2^{ts} \equiv \frac{1}{2} \left\langle \left(1-u^2\right) g_t(u) \right\rangle_{\mathcal{N}} \left\langle \left(1-u^2\right) g_s(u) \right\rangle_{\mathcal{N}} \,. \tag{31c}$$

To calculate the approximation error, $\epsilon_{apr}$ (Eq. (10)), we need $\boldsymbol{G}_s^{-1}$. For that we express $\boldsymbol{G}_s$ in terms of its eigenvalues and eigenvectors, from which the inverse follows easily. That analysis, which is nontrivial, is carried out in §8; we simply report the result here,

$$\boldsymbol{G}_s \approx \lambda^{(0)} \boldsymbol{v}^{(0)} \left(\boldsymbol{v}^{(0)}\right)^T + \sum_{k=1}^{L_1} \lambda_k^{(1)} \boldsymbol{v}_k^{(1)} \left(\boldsymbol{v}_k^{(1)}\right)^T + \sum_{k=1}^{L_2} \lambda_k^{(2)} \boldsymbol{v}_k^{(2)} \left(\boldsymbol{v}_k^{(2)}\right)^T + \sum_{k=1}^{L_r} \lambda^{(r)} \boldsymbol{v}_k^{(r)} \left(\boldsymbol{v}_k^{(r)}\right)^T \,. \tag{32}$$

As shown in §8, the eigenvalues are

$$\lambda^{(0)} = C_0^{ss} \left(c_0 + L_h\right) \tag{33a}$$

$$\lambda_k^{(1)} = C_1^{ss} \left(c_1 + \tilde{\lambda}_k^{(1)}\right) \tag{33b}$$

$$\lambda_k^{(2)} = C_2^{ss} \left(c_2 + \tilde{\lambda}_k^{(2)}\right) \tag{33c}$$

$$\lambda^{(r)} = \delta_s \,, \tag{33d}$$

where $\tilde{\lambda}_k^{(1)}$ and $\tilde{\lambda}_k^{(2)}$ are eigenvalues of $\boldsymbol{J}_s \boldsymbol{J}_s^T$ and $\boldsymbol{M}_s \boldsymbol{M}_s^T$, respectively, and the coefficients are

$$c_0 \equiv \frac{\delta_s + C_1^{ss} + C_2^{ss}}{C_0^{ss}} \tag{34a}$$

$$c_1 \equiv \frac{\delta_s + C_2^{ss}}{C_1^{ss}} \tag{34b}$$

$$c_2 \equiv \frac{\delta_s}{C_2^{ss}} \,. \tag{34c}$$

The rank of $\boldsymbol{J}_s \boldsymbol{J}_s^T$ and $\boldsymbol{M}_s \boldsymbol{M}_s^T$ are $L_1$ and $L_2$, respectively, and $L_r$ is such that it picks up any dimensionality uncaptured by the second-order expansion, (because $\boldsymbol{G}_s$ is typically full rank under a nonlinear activation function),

$$L_1 = \min[L_x, L_h - 1] \approx \min[L_x, L_h] \tag{35a}$$

$$L_2 = \min\left[\frac{1}{2} L_x(L_x-1), L_h - L_x - 1\right]^+ \approx \min\left[\frac{L_x^2}{2}, L_h - L_x\right]^+ \tag{35b}$$

$$L_r = \left[L_h - \left(1 + L_x + \frac{1}{2} L_x(L_x-1)\right)\right]^+ \approx \left[L_h - \frac{L_x^2}{2}\right]^+ \tag{35c}$$

where the superscript $+$ indicates the threshold-linear operation: $[x]^+ = x$ if $x > 0$ and 0 otherwise, and the approximations are valid because we are interested in the large $L_x$ and $L_h$ limit. The first of these two quantities, $L_1$ and $L_2$, are plotted in Fig. 5A.

We are now in a position to derive an explicit expression for the approximation error, $\epsilon_{apr}$, given in Eq. (10). Noticing that $\boldsymbol{w}_t$ is a zero-mean random vector (Eq. (2a)), in the large $L_t$ limit we may make the approximation $\boldsymbol{w}_t\boldsymbol{w}_t^T \approx \boldsymbol{I}/L_t$. Consequently,

$$\epsilon_{apr} \approx \frac{1}{L_t}\,\mathsf{Tr}\left[\langle \boldsymbol{g}_t\boldsymbol{g}_t^T\rangle - \langle \boldsymbol{g}_t\boldsymbol{g}_s^T\rangle \boldsymbol{G}_s^{-1}\langle \boldsymbol{g}_s\boldsymbol{g}_t^T\rangle\right]\,. \tag{36}$$

The first term is given by

$$\frac{1}{L_t}\,\mathsf{Tr}\left[\langle \boldsymbol{g}_t\boldsymbol{g}_t^T\rangle\right] = \int_{-\infty}^{\infty}\frac{du}{\sqrt{2\pi}}\exp\left(-\frac{u^2}{2}\right) g_t(u)^2 \equiv D_0^t\,. \tag{37}$$

To compute the second term, we start by writing it

$$\frac{1}{L_t}\,\mathsf{Tr}\left[\langle \boldsymbol{g}_t\boldsymbol{g}_s^T\rangle \boldsymbol{G}_s^{-1}\langle \boldsymbol{g}_s\boldsymbol{g}_t^T\rangle\right] = \frac{1}{L_t}\,\mathsf{Tr}\left[\langle \boldsymbol{g}_s\boldsymbol{g}_t^T\rangle\langle \boldsymbol{g}_t\boldsymbol{g}_s^T\rangle \boldsymbol{G}_s^{-1}\right]\,. \tag{38}$$

Using Eq. (29), we have

$$\begin{aligned}\frac{1}{L_t}\,\langle \boldsymbol{g}_s\boldsymbol{g}_t^T\rangle\langle \boldsymbol{g}_t\boldsymbol{g}_s^T\rangle &\approx \frac{1}{L_t}\left[C_0^{ts}\boldsymbol{1}_h\boldsymbol{1}_t^T + C_1^{ts}\boldsymbol{J}_s\boldsymbol{J}_t^T + C_2^{ts}\boldsymbol{M}_s\boldsymbol{M}_t^T\right]\left[C_0^{ts}\boldsymbol{1}_t\boldsymbol{1}_h^T + C_1^{ts}\boldsymbol{J}_t\boldsymbol{J}_s^T + C_2^{ts}\boldsymbol{M}_t\boldsymbol{M}_s^T\right]\\ &\approx (C_0^{ts})^2\boldsymbol{1}_h\boldsymbol{1}_h^T + \frac{(C_1^{ts})^2\boldsymbol{J}_s\boldsymbol{J}_s^T}{L_x} + \frac{(C_2^{ts})^2\boldsymbol{M}_s\boldsymbol{M}_s^T}{L_x^2/2}\,.\end{aligned} \tag{39}$$

The second line follows because the cross terms, $\boldsymbol{1}_h^T\boldsymbol{J}_t$, $\boldsymbol{1}_h^T\boldsymbol{M}_t$ and $\boldsymbol{J}_t^T\boldsymbol{M}_t$, are all approximately zero, so long as $L_t$ is sufficiently large. To derive this expression we took the large $L_x$ limit, and replaced $L_x(L_x-1)$ with $L_x^2$. Combining this with the expression for $\boldsymbol{G}_s$, Eq. (32), from which it is easy to write down the inverse, and making use of Eqs. (33) and (158), we arrive at

$$\frac{1}{L_t}\,\mathsf{Tr}\left[\langle \boldsymbol{g}_s\boldsymbol{g}_t^T\rangle\langle \boldsymbol{g}_t\boldsymbol{g}_s^T\rangle \boldsymbol{G}_s^{-1}\right] = \frac{(C_0^{ts})^2}{C_0^{ss}}\frac{L_h}{c_0+L_h} + \frac{1}{L_x}\frac{(C_1^{ts})^2}{C_1^{ss}}\sum_{k=1}^{L_1}\frac{\tilde{\lambda}_k^{(1)}}{c_1+\tilde{\lambda}_k^{(1)}} + \frac{1}{L_x^2/2}\frac{(C_2^{ts})^2}{C_2^{ss}}\sum_{k=1}^{L_2}\frac{\tilde{\lambda}_k^{(2)}}{c_2+\tilde{\lambda}_k^{(2)}}\,. \tag{40}$$

Given our assumption that the eigenvalue spectrum of both $\boldsymbol{J}_s\boldsymbol{J}_s^T$ and $\boldsymbol{M}_s\boldsymbol{M}_s^T$ follow the Marchenko-Pastur distribution, the sums over the eigenvalues turn into averages over the Marchenko-Pastur distribution. Those averages, which are tedious but straightforward, are computed in §9, and we arrive at

$$\begin{aligned}\frac{1}{L_t}\,\mathsf{Tr}\left[\langle \boldsymbol{g}_s\boldsymbol{g}_t^T\rangle\langle \boldsymbol{g}_t\boldsymbol{g}_s^T\rangle \boldsymbol{G}_s^{-1}\right] &= \frac{(C_0^{ts})^2}{C_0^{ss}}\left[1-\frac{c_0}{c_0+L_h}\right] + \frac{(C_1^{ts})^2}{C_1^{ss}}\left[1-f\left(\frac{L_h}{L_x};c_1\right)\right]\\ &+ \frac{(C_2^{ts})^2}{C_2^{ss}}\left[1-\frac{L_x}{L_h}\right]^+\left[1-f\left(\frac{L_h}{L_x^2/2};\frac{c_2}{1-L_x/L_h}\right)\right]\,,\end{aligned} \tag{41}$$

where $f(\bar{\lambda};c)$ is defined in Eq. (161); we repeat its definition here for convenience,

$$f(\bar{\lambda};c) \equiv \frac{\sqrt{\left(\bar{\lambda}-1+c\right)^2+4c}-\left(\bar{\lambda}-1+c\right)}{2}\,. \tag{42}$$

This function has relatively simple asymptotic behavior: it is 1 when $\bar{\lambda}=0$ and falls off as $c/\bar{\lambda}$ when $\bar{\lambda}\gg c$.

Combining Eq. (41) with the first term in the expression for the approximation error, Eq. (37), and inserting that into Eq. (10), we arrive at

$$\epsilon_{apr} \approx \delta_{ts} + \sum_{q=0}^{2}\frac{(C_q^{ts})^2}{C_q^{ss}}f_q(L_h) \tag{43}$$

where

$$f_0(L_h) \equiv \frac{c_0}{c_0+L_h} \tag{44a}$$

$$f_1(L_h) \equiv f\left(\frac{L_h}{L_x};c_1\right) \tag{44b}$$

$$f_2(L_h) \equiv \min\left[1,\frac{L_x}{L_h}\right] + \left[1-\frac{L_x}{L_h}\right]^+ f\left(\frac{L_h}{L_x^2/2};\frac{c_2}{1-L_x/L_h}\right) \approx f\left(\frac{L_h}{L_x^2/2};c_2\right) \tag{44c}$$

and

$$\delta_{ts} \equiv D_0^t - \frac{(C_0^{ts})^2}{C_0^{ss}} - \frac{(C_1^{ts})^2}{C_1^{ss}} - \frac{(C_2^{ts})^2}{C_2^{ss}} \,. \tag{45}$$

The approximation made in Eq. (44c) is accurate everywhere except the region $L_x \lesssim L_h$; that's because $f(\overline{\lambda}; c) \to 1$ when $\overline{\lambda} \ll 1$. From Eqs. (43) and (44), we recover the expression for the approximation error in the main text, with coefficients given by

$$\alpha \equiv \delta_{ts} \tag{46a}$$
$$a_0 \equiv c_0 (C_0^{ts})^2 / C_0^{ss} \tag{46b}$$
$$a_q \equiv (C_q^{ts})^2 / C_q^{ss} \,, \quad q = 1, 2 \,. \tag{46c}$$

For Eq. (46b) we used $c_0 \ll L_h$.

### 4 Estimation error

The estimation error, which is given Eq. (8b), can be written

$$\epsilon_{est} \equiv (\boldsymbol{w}_s^* - \boldsymbol{w}_s)^T \boldsymbol{G}_s (\boldsymbol{w}_s^* - \boldsymbol{w}_s). \tag{47}$$

This quantity is a random variable that depends on the data. We thus consider its mean, which is the expectation over the distribution of training data,

$$\bar{\epsilon}_{est} \equiv \left\langle (\boldsymbol{w}_s^* - \boldsymbol{w}_s)^T \boldsymbol{G}_s (\boldsymbol{w}_s^* - \boldsymbol{w}_s) \right\rangle_{p(\boldsymbol{x}_{1:N}, y_{1:N})} \tag{48}$$

where $\{\boldsymbol{x}_{1:N}, y_{1:N}\}$ is the training data. We first consider maximum likelihood, then stochastic gradient descent.

#### 4.1 Estimation error under maximum likelihood learning

As the teacher noise, $\sigma_t \xi$, is Gaussian, given $N$ training points $D_N = \{\boldsymbol{x}_n, y_n\}_{n=1}^N$ with $y_n = \boldsymbol{w}_t \cdot g(\boldsymbol{J}_t \boldsymbol{x}_n) + \sigma_t \xi_n$, the ML weights of the student network are given by the usual expression for least squares minimization,

$$\boldsymbol{w}_s = \left( \frac{1}{N} \sum_{n=1}^N g(\boldsymbol{J}_s \boldsymbol{x}_n) g(\boldsymbol{J}_s \boldsymbol{x}_n)^T \right)^{-1} \left( \frac{1}{N} \sum_{n=1}^N g(\boldsymbol{J}_s \boldsymbol{x}_n) y_n \right). \tag{49}$$

Note that $\sum_{n=1}^N g(\boldsymbol{J}_s \boldsymbol{x}_n) g(\boldsymbol{J}_s \boldsymbol{x}_n)^T$ is not invertible unless $N > L_h$, so we work in that regime. Denoting

$$\boldsymbol{g}_t^n \equiv g(\boldsymbol{J}_t \boldsymbol{x}_n) \tag{50a}$$
$$\boldsymbol{g}_s^n \equiv g(\boldsymbol{J}_s \boldsymbol{x}_n) \tag{50b}$$
$$\boldsymbol{G}_s^{(N)} \equiv \frac{1}{N} \sum_{n=1}^N \boldsymbol{g}_s^n \left( \boldsymbol{g}_s^n \right)^T \tag{50c}$$

and noting that $y_n = \boldsymbol{w}_t \cdot \boldsymbol{g}_t^n + \sigma_t \xi_n$, $\boldsymbol{w}_s - \boldsymbol{w}_s^*$ is written

$$\boldsymbol{w}_s - \boldsymbol{w}_s^* = \left( \boldsymbol{G}_s^{(N)} \right)^{-1} \left( \frac{1}{N} \sum_{n=1}^N \boldsymbol{g}_s^n \left[ \boldsymbol{w}_t \cdot \boldsymbol{g}_t^n + \sigma_t \xi_n - \boldsymbol{w}_s^* \cdot \boldsymbol{g}_s^n \right] \right). \tag{51}$$

Inserting this into Eq. (47), we have

$$\bar{\epsilon}_{est} = \frac{1}{N^2} \Bigg\langle \left( \sum_{n=1}^N \left[ \boldsymbol{w}_t \cdot \boldsymbol{g}_t^n + \sigma_t \xi_n - \boldsymbol{w}_s^* \cdot \boldsymbol{g}_s^n \right] (\boldsymbol{g}_s^n)^T \right) \left( \boldsymbol{G}_s^{(N)} \right)^{-1} \boldsymbol{G}_s \left( \boldsymbol{G}_s^{(N)} \right)^{-1}$$
$$\left( \sum_{n'=1}^N \boldsymbol{g}_s^{n'} \left[ \boldsymbol{w}_t \cdot \boldsymbol{g}_t^{n'} + \sigma_t \xi_{n'} - \boldsymbol{w}_s^* \cdot \boldsymbol{g}_s^{n'} \right] \right) \Bigg\rangle. \tag{52}$$

The first observation is that the $n$ and $n'$-dependent terms are independent when $n \neq n'$. Consequently, the double sum over $n$ and $n'$ can be replaced by its diagonal elements,

$$\bar{\epsilon}_{est} \approx \frac{1}{N^2} \left\langle \sum_{n=1}^{N} \left[\boldsymbol{w}_t \cdot \boldsymbol{g}_t^n + \sigma_t \xi_n - \boldsymbol{w}_s^* \cdot \boldsymbol{g}_s^n\right]^2 (\boldsymbol{g}_s^n)^T \left(\boldsymbol{G}_s^{(N)}\right)^{-1} \boldsymbol{G}_s \left(\boldsymbol{G}_s^{(N)}\right)^{-1} \boldsymbol{g}_s^n \right\rangle . \tag{53}$$

Second, we assume that $[\boldsymbol{w}_t \cdot \boldsymbol{g}_t^n - \boldsymbol{w}_s^* \cdot \boldsymbol{g}_s^n]^2$ and $(\boldsymbol{g}_s^n)^T \boldsymbol{g}_s^n$ average independently. Using Eqs. (8a) and (50c), this leads to

$$\begin{aligned} \bar{\epsilon}_{est} &\approx \left\langle [\boldsymbol{w}_t \cdot \boldsymbol{g}_t + \sigma_t \xi - \boldsymbol{w}_s^* \cdot \boldsymbol{g}_s]^2 \right\rangle \frac{1}{N} \left\langle \mathsf{Tr} \left[ \left(\boldsymbol{G}_s^{(N)}\right)^{-1} \boldsymbol{G}_s \left(\boldsymbol{G}_s^{(N)}\right)^{-1} \frac{1}{N} \sum_{n=1}^{N} \boldsymbol{g}_s^n (\boldsymbol{g}_s^n)^T \right] \right\rangle \\ &= \left(\epsilon_{apr} + \sigma_t^2\right) \frac{1}{N} \mathsf{Tr} \left[ \left(\boldsymbol{G}_s^{(N)}\right)^{-1} \boldsymbol{G}_s \right] . \end{aligned} \tag{54}$$

To derive an explicit expression for $\bar{\epsilon}_{est}$, we need to determine how the trace term scales as $N$ and $L_h$ go to infinity, with the ratio $L_h/N$ fixed at some value less than 1. We start by turning $\boldsymbol{G}_s^{(N)}$ into a zero mean matrix. To that end, we let $\boldsymbol{G}_s^{(N)} \equiv \delta \boldsymbol{G}_s^{(N)} + \bar{\boldsymbol{g}}_N \bar{\boldsymbol{g}}_N^T$ where

$$\delta \boldsymbol{G}_s^{(N)} \equiv \frac{1}{N} \sum_{n=1}^{N} (\boldsymbol{g}_s^n - \bar{\boldsymbol{g}}_n)(\boldsymbol{g}_s^n - \bar{\boldsymbol{g}}_n)^T \tag{55}$$

and

$$\bar{\boldsymbol{g}}_N \equiv \frac{1}{N} \sum_{n=1}^{N} \boldsymbol{g}_s^n \approx \langle \boldsymbol{g}_s \rangle_{p(\boldsymbol{x})} \equiv \bar{\boldsymbol{g}} . \tag{56}$$

Further approximating $\boldsymbol{G}_s^{(N)}$ with $\delta \boldsymbol{G}_s^{(N)} + \bar{\boldsymbol{g}} \bar{\boldsymbol{g}}^T$, and applying the Sherman-Morrison formula, we have

$$\left(\boldsymbol{G}_s^{(N)}\right)^{-1} \approx \left(\delta \boldsymbol{G}_s^{(N)} + \bar{\boldsymbol{g}} \bar{\boldsymbol{g}}^T\right)^{-1} = \left(\delta \boldsymbol{G}_s^{(N)}\right)^{-1} - \frac{\left(\delta \boldsymbol{G}_s^{(N)}\right)^{-1} \bar{\boldsymbol{g}} \bar{\boldsymbol{g}}^T \left(\delta \boldsymbol{G}_s^{(N)}\right)^{-1}}{1 + \bar{\boldsymbol{g}}^T \left(\delta \boldsymbol{G}_s^{(N)}\right)^{-1} \bar{\boldsymbol{g}}} . \tag{57}$$

Next we need an approximation for $\boldsymbol{G}_s$. In §3, we used Eq. (28). Here we make a more severe aproximation: decomposing $\boldsymbol{G}_s$ as $\delta \boldsymbol{G}_s + \bar{\boldsymbol{g}} \bar{\boldsymbol{g}}^T$, we use $\delta \boldsymbol{G}_s \approx \sigma_M^2 \boldsymbol{I}$. This is consistent with approximating $\delta \boldsymbol{G}_s^{(N)}$ as a Wishart matrix $W_{L_h}(\frac{\sigma_M^2}{N} \boldsymbol{I}, N)$, which converges to $\sigma_M^2 \boldsymbol{I}$ as $N \to \infty$. Inserting this, along with the above expression for $\boldsymbol{G}_s^{(N)}$, into Eq. (54), we arrive, after a small amount of algebra, at

$$\mathsf{Tr} \left[ \left(\boldsymbol{G}_s^{(N)}\right)^{-1} \boldsymbol{G}_s \right] \approx \sigma_M^2 \mathsf{Tr} \left[ \left(\delta \boldsymbol{G}_s^{(N)}\right)^{-1} \right] + 1 - \frac{1 + \sigma_M^2 \bar{\boldsymbol{g}}^T \left(\delta \boldsymbol{G}_s^{(N)}\right)^{-2} \bar{\boldsymbol{g}}}{1 + \bar{\boldsymbol{g}}^T \left(\delta \boldsymbol{G}_s^{(N)}\right)^{-1} \bar{\boldsymbol{g}}} . \tag{58}$$

The first term is proportional to $L_h$, the dimensionality of $\delta \boldsymbol{G}_s^{(N)}$. The last is bounded by $(1 + \sigma_M^2 |\bar{\boldsymbol{g}}|^2 / \lambda_{\min}^2)/(1 + |\bar{\boldsymbol{g}}|^2 / \lambda_{\max})$ where $\lambda_{\min}$ and $\lambda_{\max}$ are the minimum and maximum eigenvalues of $\delta \boldsymbol{G}_s^{(N)}$. Since $\delta \boldsymbol{G}_s^{(N)}$ is the sum of $N$ outer products of random vectors, its spectrum approximately follows a Marchenko-Pastur distribution with parameters $\sigma_M^2$ and $L_h/N$. Assuming $L_h$ is not too close to $N$, the eigenvalues are $\mathcal{O}(1)$. Consequently, the above expression is dominated by the first term. Restoring the prefactor $1/N$, we have

$$\frac{1}{N} \mathsf{Tr} \left[ \left(\boldsymbol{G}_s^{(N)}\right)^{-1} \boldsymbol{G}_s \right] \approx \frac{\sigma_M^2 L_h}{N} \left\langle \frac{1}{\lambda} \right\rangle_{MP(\sigma_M^2, L_h/N)} = \frac{L_h}{N - L_h} \tag{59}$$

where the average over $\lambda^{-1}$ (which gives us the second equality) is computed in §9 (see in particular Eq. (167)).

Equation (59) is consistent with previous work on linear regression using the replica method [24]. Inserting Eq. (59) into (54), we arrive at

$$\bar{\epsilon}_{est} \approx (\epsilon_{apr} + \sigma_t^2) \frac{L_h}{N - L_h} . \tag{60}$$

Consequently, the generalization error, $\epsilon_{gen} = \bar{\epsilon}_{est} + \epsilon_{apr} + \sigma_t^2$, Eq. (7), is given approximately by

$$\epsilon_{gen} \approx (\epsilon_{apr} + \sigma_t^2) \frac{N}{N - L_h} . \tag{61}$$

#### 4.2 Estimation error under stochastic gradient descent (SGD) learning

In an online setting, it is more realistic to consider stochastic gradient descent rather than maximum likelihood, the former given by

$$\boldsymbol{w}_s^{(n)} = \boldsymbol{w}_s^{(n-1)} + \eta(y_n - \hat{y}_n)\boldsymbol{g}_s^n \tag{62}$$

where $\boldsymbol{g}_s^n$ is defined in Eq. (50b) and $\hat{y}_n = \boldsymbol{w}_s^{(n-1)} \cdot \boldsymbol{g}_s^n$ (Eq. (3)). Making the definition

$$\boldsymbol{u}_n \equiv \boldsymbol{w}_s^{(n)} - \boldsymbol{w}_s^*, \tag{63}$$

the update rule for $\boldsymbol{u}_n$ is

$$\begin{aligned}\boldsymbol{u}_n &= \left[\boldsymbol{I} - \eta \boldsymbol{g}_s^n (\boldsymbol{g}_s^n)^T\right] \boldsymbol{u}_{n-1} + \eta \left[\boldsymbol{w}_t \cdot \boldsymbol{g}_t^n - \boldsymbol{w}_s^* \cdot \boldsymbol{g}_s^n + \sigma_t \xi_n\right] \boldsymbol{g}_s^n \\ &= \left[\boldsymbol{I} - \eta \boldsymbol{G}_s\right] \boldsymbol{u}_{n-1} + \eta \left[\boldsymbol{G}_s - \boldsymbol{g}_s^n (\boldsymbol{g}_s^n)^T\right] \boldsymbol{u}_{n-1} + \eta \left[\boldsymbol{w}_t \cdot \boldsymbol{g}_t^n - \boldsymbol{w}_s^* \cdot \boldsymbol{g}_s^n + \sigma_t \xi_n\right] \boldsymbol{g}_s^n\end{aligned} \tag{64}$$

where, recall, $\boldsymbol{G}_s$ is the hidden layer covariance, defined in Eq. (11). After $n$ updates, the estimation error (Eq. (8b)) is

$$\epsilon_{est}^{(n)} = \boldsymbol{u}_n^T \boldsymbol{G}_s \boldsymbol{u}_n. \tag{65}$$

It is convenient to work in a basis spanned by the eigenvectors of $\boldsymbol{G}_s$. That basis is given in Eq. (32), which shows a great deal of structure, and in particular a division into four components. Later we will use that structure, but for now we adopt a notation that hides it: we simply write $\boldsymbol{v}_\mu$ and $\lambda_\mu$ for the $\mu^{\text{th}}$ eigenvector and eigenvalue of $\boldsymbol{G}_s$ (i.e., $\boldsymbol{G}_s \boldsymbol{v}_\mu = \lambda_\mu \boldsymbol{v}_\mu$). We then make the change of variables

$$\boldsymbol{u}_n = \sum_\mu m_{\mu,n} \boldsymbol{v}_\mu \,. \tag{66}$$

In the new variables, the estimation error is

$$\epsilon_{est}^{(n)} = \sum_\mu \lambda_\mu m_{\mu,n}^2 \,. \tag{67}$$

We'll first find the update rules for $m_{\mu,n}$, then use them to find the update rules for $m_{\mu,n}^2$. Taking the eigenvectors to be orthonormal, we have $m_{\mu,n} = \boldsymbol{v}_\mu \cdot \boldsymbol{u}_n$; applying this to Eq. (64) yields

$$m_{\mu,n} = (1 - \eta\lambda_\mu) m_{\mu,n-1} + \eta \boldsymbol{v}_\mu^T \left[\boldsymbol{G}_s - \boldsymbol{g}_s^n (\boldsymbol{g}_s^n)^T\right] \boldsymbol{u}_{n-1} + \eta \left[\boldsymbol{w}_t \cdot \boldsymbol{g}_t^n - \boldsymbol{w}_s^* \cdot \boldsymbol{g}_s^n + \sigma_t \xi_n\right] \boldsymbol{v}_\mu \cdot \boldsymbol{g}_s^n \,. \tag{68}$$

Squaring both sides gives us an expression for $m_{\mu,n}^2$. To simplify that expression, we assume that the mean dynamics of $m_{\mu,n}^2$ is described by the dynamics of the mean, $\langle m_{\mu,n}^2 \rangle$, where the average is over the distribution of the input $\boldsymbol{x}$ and the teacher noise $\xi$. To simplify notation, below we suppress the label $n$ that appears on $\boldsymbol{g}_s^n, \boldsymbol{g}_t^n$, and $\xi_n$. (Note that $\boldsymbol{g}_s^n$ and $\boldsymbol{g}_t^n$ depend on $n$ only through $\boldsymbol{x}_n$; see Eqs. (50a) and (50b)). The first term on the right hand side of Eq. (68) is independent of $\boldsymbol{x}$, and the second two terms, which do depend on $\boldsymbol{x}$, are both zero mean. We assume those terms are uncorrelated, so we have

$$\begin{aligned}m_{\mu,n}^2 = (1 - \eta\lambda_\mu)^2 m_{\mu,n-1}^2 &+ \eta^2 \left\langle \boldsymbol{u}_{n-1}^T [\boldsymbol{G}_s - \boldsymbol{g}_s \boldsymbol{g}_s^T] \boldsymbol{v}_\mu \boldsymbol{v}_\mu^T [\boldsymbol{G}_s - \boldsymbol{g}_s \boldsymbol{g}_s^T] \boldsymbol{u}_{n-1} \right\rangle \\ &+ \eta^2 \left\langle \left[\boldsymbol{w}_t \cdot \boldsymbol{g}_t - \boldsymbol{w}_s^* \cdot \boldsymbol{g}_s + \sigma_t \xi\right]^2 (\boldsymbol{v}_\mu \cdot \boldsymbol{g}_s)^2 \right\rangle \,.\end{aligned} \tag{69}$$

To simplify the first average, we note that it can be written

$$\langle \boldsymbol{u}_{n-1}^T [\boldsymbol{G}_s - \boldsymbol{g}_s \boldsymbol{g}_s^T] \boldsymbol{v}_\mu \boldsymbol{v}_\mu^T [\boldsymbol{G}_s - \boldsymbol{g}_s \boldsymbol{g}_s^T] \boldsymbol{u}_{n-1} \rangle = \langle (\boldsymbol{u}_{n-1} \cdot \boldsymbol{g}_s)^2 (\boldsymbol{v}_\mu \cdot \boldsymbol{g}_s)^2 \rangle - \left(\boldsymbol{u}_{n-1}^T \boldsymbol{G}_s \boldsymbol{v}_\mu\right)^2 \,. \tag{70}$$

Assuming $(\boldsymbol{u}_{n-1} \cdot \boldsymbol{g}_s)^2$ and $(\boldsymbol{v}_\mu \cdot \boldsymbol{g}_s)^2$ self average, for the first term we have

$$\left\langle (\boldsymbol{u}_{n-1} \cdot \boldsymbol{g}_s)^2 (\boldsymbol{v}_\mu \cdot \boldsymbol{g}_s)^2 \right\rangle = \boldsymbol{u}_{n-1}^T \boldsymbol{G}_s \boldsymbol{u}_{n-1} \, \boldsymbol{v}_\mu^T \boldsymbol{G}_s \boldsymbol{v}_\mu = \boldsymbol{u}_{n-1}^T \boldsymbol{G}_s \boldsymbol{u}_{n-1} \, \lambda_\mu \,. \tag{71}$$

Then, using the fact that $\boldsymbol{G}_s \boldsymbol{v}_\mu = \lambda_\mu \boldsymbol{v}_\mu$ and $\boldsymbol{u}_{n-1} \cdot \boldsymbol{v}_\mu = m_{\mu,n-1}$, we arrive at

$$\left\langle \boldsymbol{u}_{n-1}^T [\boldsymbol{G}_s - \boldsymbol{g}_s \boldsymbol{g}_s^T] \boldsymbol{v}_\mu \boldsymbol{v}_\mu^T [\boldsymbol{G}_s - \boldsymbol{g}_s \boldsymbol{g}_s^T] \boldsymbol{u}_{n-1} \right\rangle = \lambda_\mu \sum_\nu \lambda_\nu m_{\nu,n-1}^2 - \lambda_\mu^2 m_{\mu,n-1}^2 \,. \tag{72}$$

For the second average in Eq. (69), we again assume that $(\boldsymbol{v}_\mu \cdot \boldsymbol{g}_s)^2$ self averages, so the average of the product is just the product of the averages. The average of the square of the term in brackets is $\epsilon_{apr} + \sigma_t^2$ (see Eq. (8a)) and the average of $(\boldsymbol{v}_\mu \cdot \boldsymbol{g}_s)^2$ is, as in Eq. (71), $\lambda_\mu$. Thus, the second average in Eq. (69) simplifies to

$$\left\langle \left[\boldsymbol{w}_t \cdot \boldsymbol{g}_t^n - \boldsymbol{w}_s^* \cdot \boldsymbol{g}_s^n + \sigma_t \xi\right]^2 (\boldsymbol{v}_\mu \cdot \boldsymbol{g}_s)^2 \right\rangle = (\epsilon_{apr} + \sigma_t^2) \lambda_\mu \,. \tag{73}$$

Inserting Eqs. (72) and (73) into (69), we arrive at

$$m_{\mu,n}^2 = (1 - 2\eta\lambda_\mu) m_{\mu,n-1}^2 + \eta^2 \lambda_\mu \sum_\nu \lambda_\nu m_{\nu,n-1}^2 + \eta^2(\epsilon_{apr} + \sigma_t^2)\lambda_\mu \, . \tag{74}$$

To solve this equation, we define the matrix

$$A_{\mu\nu} \equiv 2\eta\lambda_\mu \delta_{\mu\nu} - \eta^2 \lambda_\mu \lambda_\nu \, . \tag{75}$$

After a small amount of algebra, we find that

$$\boldsymbol{m}_n^2 = \frac{\eta(\epsilon_{apr} + \sigma_t^2)}{2 - \eta D_0^s L_h} \mathbf{1}_h + (\boldsymbol{I} - \boldsymbol{A})^n \left( \boldsymbol{m}_0^2 - \frac{\eta(\epsilon_{apr} + \sigma_t^2)}{2 - \eta D_0^s L_h} \mathbf{1}_h \right) \tag{76}$$

where we used

$$\sum_\mu \lambda_\mu = D_0^s L_h \tag{77}$$

(which follows from Eq. (15)), $\boldsymbol{m}_n^2$ is a vector whose $\mu^{\text{th}}$ component is $m_{\mu,n}^2$, and we used the fact that $\boldsymbol{A}^{-1} \cdot \boldsymbol{\lambda} = \mathbf{1}_h/(2\eta - \eta^2 D_0^s L_h)$ where $\boldsymbol{\lambda} \equiv (\lambda_1, \lambda_2, ...)$, which follows from the Sherman-Morrison formula.

The term $(\boldsymbol{I} - \boldsymbol{A})^n$ is problematic, as its eigenvalues and eigenvectors cannot be found analytically. We thus make a very severe approximation: we let

$$A_{\mu\nu} \approx (2\eta - \eta^2 D_0^s L_h)\lambda_\mu \delta_{\mu\nu} \, . \tag{78}$$

With this approximation, Eq. (76) simplifies to

$$m_{\mu,n}^2 = \frac{\eta(\epsilon_{apr} + \sigma_t^2)}{2 - \eta D_0^s L_h} + \left(1 - \eta\lambda_\mu(2 - \eta D_0^s L_h)\right)^n \left( m_{\mu,0}^2 - \frac{\eta(\epsilon_{apr} + \sigma_t^2)}{2 - \eta D_0^s L_h} \right) \, . \tag{79}$$

We choose $\eta$ to maximize the rate of decay of $m_{\mu,n}^2$ (that is, minimize $1 - \eta\lambda_\mu(2 - \eta D_0^s L_h)$); this yields

$$\eta^* = \frac{1}{D_0^s L_h} \, . \tag{80}$$

All the eigenmodes show the fastest decay at this learning rate regardless of their eigenvalues. Replacing $\eta$ with $\eta^*$ in Eq. (79), we have

$$m_{\mu,n}^2 = m_\infty^2 + \left( m_{\mu,0}^2 - m_\infty^2 \right) \left( 1 - \frac{\lambda_\mu}{D_0^s L_h} \right)^n \, . \tag{81}$$

where

$$m_\infty^2 \equiv \frac{\epsilon_{apr} + \sigma_t^2}{D_0^s L_h} \, . \tag{82}$$

We can use this expression to determine how the estimation error, Eq. (67) evolves in time. Inserting Eq. (81) into (67), the average estimation error after $n$ samples, $\bar{\epsilon}_{est}^{(n)}$, is given by

$$\bar{\epsilon}_{est}^{(n)} = \sum_\mu \left( \lambda_\mu m_\infty^2 + \lambda_\mu (m_{\mu,0}^2 - m_\infty^2) \left[ 1 - \frac{\lambda_\mu}{D_0^s L_h} \right]^n \right) \, . \tag{83}$$

We now take advantage of the structure implicit in Eq. (32), which tells us that the eigenvalues are divided into four components, which we label with $q \in \{0, 1, 2, r\}$. We thus have

$$\bar{\epsilon}_{est}^{(n)} = \sum_{q \in \{0,1,2,r\}} \sum_{\mu \in S_q} \left( \lambda_\mu m_\infty^2 + \lambda_\mu (m_{\mu,0}^2 - m_\infty^2) \left[ 1 - \frac{\lambda_\mu}{D_0^s L_h} \right]^n \right) \tag{84}$$

where $S_q$ specifies the range of $\mu$,

$$S_0\colon\ \mu = 1 \tag{85a}$$

$$S_1\colon\ 1 < \mu \leq L_1 + 1 \tag{85b}$$

$$S_2\colon\ L_1 + 1 < \mu \leq L_2 + L_1 + 1 \tag{85c}$$

$$S_r\colon\ L_2 + L_1 + 1 < \mu \leq L_h \, . \tag{85d}$$

The first component, $S_0$, contains only one eigenmode, which corresponds to the largest eigenvalue $\lambda_1$ ($= \lambda^{(0)}$ in Eq. (33)). The rest contain multiple eigenmodes. For those modes we can approximate the exponential term as

$$(1 - \lambda_\mu/(D_0^s L_h))^n \approx e^{-n\lambda_\mu/(D_0^s L_h)} \approx e^{-n\langle\lambda_\mu\rangle_q/(D_0^s L_h)} \tag{86}$$

where the subscript $q$ means an average over $\mu \in S_q$. For the first inequality we used $\lambda_\mu \ll L_h$ for $\mu > 1$; for the second we used that fact that the eigenvalues typically have a small spread within each component. Making that replacement in Eq. (84), the lifetime cumulative estimation error, denoted $\bar{\epsilon}_{cml}^N$, is given by

$$\bar{\epsilon}_{cml}^N \equiv \frac{1}{N}\sum_{n=0}^{N-1} \bar{\epsilon}_{est}^n = \sum_{q\in\{0,1,2,r\}} \sum_{\mu\in S_q} \left(\lambda_\mu m_\infty^2 + \lambda_\mu (m_{\mu,0}^2 - m_\infty^2) R_q(L_h)\right) \tag{87}$$

where

$$R_q(L_h) \equiv \begin{cases} \frac{D_0^s L_h}{N\lambda^{(0)}}\left[1 - \left(1 - \frac{\lambda^{(0)}}{D_0^s L_h}\right)^N\right] & q = 0 \\ \frac{D_0^s L_h}{N\langle\lambda_\mu\rangle_q}\left[1 - e^{-N\langle\lambda_\mu\rangle_q/(D_0^s L_h)}\right] & \text{otherwise} \end{cases} . \tag{88}$$

The function $R_q(L_h)$ scales as $D_0^s L_h/N\langle\lambda_\mu\rangle_q$ when $L_h \ll N\langle\lambda_\mu\rangle_q$ and approaches 1 when $L_h \gg N\langle\lambda_\mu\rangle_q$.

In §8 we computed the average eigenvalues (see Eq. (157)), so the only quantity we do not know is the average over $\lambda_\mu m_{\mu,0}^2$. That quantity is computed in the next section; using that result and applying a small amount of algebra, we arrive at

$$\bar{\epsilon}_{cml}^N = \sum_q \left( L_q \langle\lambda_\mu\rangle_q \left[ m_\infty^2 (1 - R_q(L_h)) + \frac{\sigma_R^2}{L_h} R_q(L_h) \right] + \frac{(C_q^{ts})^2}{C_q^{ss}} (1 - f_q(L_h)) R_q(L_h) \right) \tag{89}$$

where

$$C_r^{ts} \equiv 0\,, \tag{90a}$$

$$L_0 \equiv 1\,, \tag{90b}$$

$\sigma_R^2/L_h$ is the initial variance of the weights, and $f_0$, $f_1$ and $f_2$ are defined in Eq. (44). (Because $C_r^{ts} = 0$, we do not need to define $f_r$.)

In Eq. (15) of the main text, we write down an expression for the average estimation error versus $n$. Here we derive that expression. As can be seen by comparing Eqs. (84) and (87) and taking into account the approximation made in Eq. (86), the only difference between $\bar{\epsilon}_{est}^{(n)}$ and $\bar{\epsilon}_{cml}^N$ is that $(1 - \langle\lambda_\mu\rangle_q/D_0^s L_h)^n$ is replaced by $R_q(L_h)$. Making the reverse replacement in Eq. (87), and approximating $(1 - \langle\lambda_\mu\rangle_q/D_0^s L_h)^n$ by $e^{-n\langle\lambda_\mu\rangle_q/D_0^s L_h}$, we have

$$\bar{\epsilon}_{est}^n = \epsilon_{apr} + \sigma_t^2 + \sum_q \left[ \frac{L_q\langle\lambda_\mu\rangle_q}{D_0^s L_h}\left(D_0^s\sigma_R^2 - (\epsilon_{apr} + \sigma_t^2)\right) + \frac{(C_q^{ts})^2}{C_q^{ss}}(1 - f_q(L_h)) \right] e^{-n\langle\lambda_\mu\rangle_q/(D_0^s L_h)} . \tag{91}$$

To derive this expression, we used the fact that $\sum_q L_q\langle\lambda_\mu\rangle_q = \sum_\mu \lambda_\mu = D_0^s L_h$ (see Eq. (77) for the second inequality), and we replaced $m_\infty^2$ by $(\epsilon_{apr} + \sigma_t^2)/D_0^s L_h$ (see Eq. (82)). The terms in square brackets correspond to the $b_q$ in Eq. (15) of the main text. The terms in the exponents were approximated from Eq. (35) and Eq. (157) as

$$\frac{\langle\lambda_\mu\rangle_q}{D_0^s L_h} \approx \frac{C_q^{ss}}{D_0^s L_q}\,, \tag{92}$$

where the expression for $q = 2$ is valid in the regime $L_h \gg L_x$. For $D_0^s$ and $C_q^{ss}$ we used Eqs. (136) and (137), respectively.

**Initial conditions**

To estimate the contribution from the initial conditions (the term containing $\lambda_\mu m_{\mu,0}^2$ in Eq. (87)), we need an expression for $m_{\mu,0}$. Using $m_{\mu,0} = \boldsymbol{v}_\mu \cdot \boldsymbol{u}_0$, we write

$$m_{\mu,0}^2 = \left(\boldsymbol{v}_\mu \cdot [\boldsymbol{w}_s^{(0)} - \boldsymbol{w}_s^*]\right)^2 \approx \left(\boldsymbol{v}_\mu \cdot \boldsymbol{w}_s^{(0)}\right)^2 + (\boldsymbol{v}_\mu \cdot \boldsymbol{w}_s^*)^2 . \tag{93}$$

The projection weights are initialized as $\boldsymbol{w}_s^{(0)} \sim N(0, \sigma_R^2/L_h)$, so the first term is given approximately by

$$\left(\boldsymbol{v}_\mu \cdot \boldsymbol{w}_s^{(0)}\right)^2 \approx \sum_{j=1}^{L_h} (v_{\mu,j})^2 \left(\boldsymbol{w}_{s,j}^{(0)}\right)^2 \approx \frac{\sigma_R^2}{L_h}\,. \tag{94}$$

For the second term we use Eq. (6) for $\boldsymbol{w}^*$, leading to

$$(\boldsymbol{v}_\mu \cdot \boldsymbol{w}_s^*)^2 = \boldsymbol{v}_\mu^T \boldsymbol{G}_s^{-1} \langle \boldsymbol{g}_s \boldsymbol{g}_t^T \rangle \boldsymbol{w}_t \boldsymbol{w}_t^T \langle \boldsymbol{g}_t \boldsymbol{g}_s^T \rangle \boldsymbol{G}_s^{-1} \boldsymbol{v}_\mu \approx \frac{1}{L_t} \mathrm{Tr}\left[\boldsymbol{v}_\mu \boldsymbol{v}_\mu^T \boldsymbol{G}_s^{-1} \langle \boldsymbol{g}_s \boldsymbol{g}_t^T \rangle \langle \boldsymbol{g}_t \boldsymbol{g}_s^T \rangle \boldsymbol{G}_s^{-1}\right] \tag{95}$$

where the approximate expression follows from $\boldsymbol{w}_t \boldsymbol{w}_t^T \approx \boldsymbol{I}/L_t$. If we were to multiply the right hand side by $\lambda_\mu$ and sum over all $\mu$, we would recover the left hand side of Eq. (41), because $\sum_\mu \lambda_\mu \boldsymbol{v}_\mu \boldsymbol{v}_\mu^T = \boldsymbol{G}_s$. Therefore, we can read off the sum of each component of $\mu$ from the right hand side of Eq. (41),

$$\sum_{\mu \in S_0} \lambda_\mu \left(\boldsymbol{v}_\mu \cdot \boldsymbol{w}_s^*\right)^2 \approx \frac{(C_0^{ts})^2}{C_0^{ss}} \left[1 - \frac{c_0}{c_0 + L_h}\right] \tag{96a}$$

$$\sum_{\mu \in S_1} \lambda_\mu \left(\boldsymbol{v}_\mu \cdot \boldsymbol{w}_s^*\right)^2 \approx \frac{(C_1^{ts})^2}{C_1^{ss}} \left[1 - f\left(\frac{L_h}{L_x}; c_1\right)\right] \tag{96b}$$

$$\sum_{\mu \in S_2} \lambda_\mu \left(\boldsymbol{v}_\mu \cdot \boldsymbol{w}_s^*\right)^2 \approx \frac{(C_2^{ts})^2}{C_2^{ss}} \left[1 - \frac{L_x}{L_h}\right]^+ \left[1 - f\left(\frac{L_h}{L_x^2/2}; \frac{c_2}{1 - L_x/L_h}\right)\right] \tag{96c}$$

$$\sum_{\mu \in S_r} \lambda_\mu \left(\boldsymbol{v}_\mu \cdot \boldsymbol{w}_s^*\right)^2 \approx 0. \tag{96d}$$

### 5 Generalization error

To determine how the optimal hidden layer size, denoted $L_h^*$, scales with the input layer size, $L_x$, we need to minimize the generalization error (found by combining the approximation and estimation errors; see Eq. (7)) with respect to $L_h$. This is nontrivial: as can be seen in Figs. 4A and 5D, the optimum exhibits three different regimes, depending on the input layer size, $L_x$. We can, though, access these regimes by considering different relative scaling of $L_h$ and $L_x$: $L_h \gg L_x^2$, $L_x^2 \gg L_h \gg L_x$, and $L_x \gg L_h$. We begin by providing estimates for the approximation error, Eq. (43), in the three regimes; in the next two sections we use those results to compute the generalization error, first for maximum likelihood learning and then for stochastic gradient descent.

To see how the approximation error, Eq. (43), scales with $L_h$, note that (as mentioned after Eq. (42)) $f(\bar{\lambda}; c) \to c/\bar{\lambda}$ when $\bar{\lambda} \gg c$, and $f(\bar{\lambda}; c) \to 1$ when $\bar{\lambda} \to 0$. Using this, and the definitions of $c_0$, $c_1$ and $c_2$ in Eq. (34), it is straightforward to show that

$$\epsilon_{apr} + \sigma_t^2 \approx \begin{cases} \sigma_t^2 + \delta_{ts} + \left(\frac{C_2^{ts}}{C_2^{ss}}\right)^2 \frac{\delta_s L_x^2}{2L_h} & \text{if } L_h \gg L_x^2 \\ \sigma_t^2 + \delta_{ts} + \frac{(C_2^{ts})^2}{C_2^{ss}} + \left(\frac{C_1^{ts}}{C_1^{ss}}\right)^2 \frac{(\delta_s + C_2^{ss}) L_x}{L_h} & \text{if } L_x^2 \gg L_h \gg L_x \\ \sigma_t^2 + \delta_{ts} + \frac{(C_2^{ts})^2}{C_2^{ss}} + \frac{(C_1^{ts})^2}{C_1^{ss}} + \left(\frac{C_0^{ts}}{C_0^{ss}}\right)^2 \frac{\delta_s + C_1^{ss} + C_2^{ss}}{c_0 + L_h} & \text{if } L_x \gg L_h\,. \end{cases} \tag{97}$$

We now use these expressions to compute the hidden layer size that optimizes the generalization error, first for maximum likelihood, and then for stochastic gradient descent.

#### 5.1 Maximum likelihood

For maximum likelihood learning, the generalization error is given in Eq. (61). Our goal now is to combine that expression with Eq. (97), the approximation error, to get the generalization error, and minimize that with respect to $L_h$ to find the optimal hidden layer size. Given the complexity of the generalization error, it is not possible to perform the exact minimization analytically. However, the generalization error becomes tractable in three regimes, $L_h \gg L_x^2$, $L_x^2 \gg L_h \gg L_x$ and $L_x \gg L_h$. We thus take the following four-step approach. In step 1, we assume that $L_h$ lies in one of the regimes, say $L_h \ll L_x^2$ for definiteness. In step 2, we write down a simplified expression for the generalization error that is valid in this regime. In step 3, we find the value of $L_h$ that minimizes the (simplified) generalization error. In step 4, we ask whether the minimum lies in the relevant region, in this case $L_h \ll L_x^2$. If it does, we have found a self-consistent minimum.

###### Optimal hidden layer size when $L_h \gg L_x^2$

In this regime, the generalization error is given by

$$\epsilon_{gen} \approx \left(\sigma_t^2 + \delta_{ts} + \left(\frac{C_2^{ts}}{C_2^{ss}}\right)^2 \frac{\delta_s L_x^2}{2L_h}\right) \frac{N}{N - L_h}. \tag{98}$$

Minimizing with respect to $L_h$ yields

$$L_h^* = \sqrt{\left(B_2^{ml} L_x^2\right)^2 + B_2^{ml} N L_x^2} - B_2^{ml} L_x^2 \tag{99}$$

where

$$B_2^{ml} \equiv \left(\frac{C_2^{ts}}{C_2^{ss}}\right)^2 \frac{\delta_s}{2(\sigma_t^2 + \delta_{ts})}. \tag{100}$$

For this solution to be consistent with the condition $L_h \gg L_x^2$, $N$ must satisfy $N \gg L_x^2 / B_2^{ml}$. Therefore, if $L_x \ll \sqrt{B_2^{ml} N}$, then the hidden layer size that minimizes the generalization error is

$$L_h^* \approx \sqrt{B_2^{ml} N L_x^2}\,. \tag{101}$$

###### Optimal hidden layer size when $L_x^2 \gg L_h \gg L_x$

In this regime, the generalization error is given by

$$\epsilon_{gen} \approx \left(\sigma_t^2 + \delta_{ts} + \frac{(C_2^{ts})^2}{C_2^{ss}} + \left(\frac{C_1^{ts}}{C_1^{ss}}\right)^2 \frac{(\delta_s + C_2^{ss}) L_x}{L_h}\right) \frac{N}{N - L_h}\,. \tag{102}$$

Minimizing with respect to $L_h$ yields

$$L_h^* = \sqrt{(B_1^{ml} L_x)^2 + B_1^{ml} N L_x} - B_1^{ml} L_x \tag{103}$$

where

$$B_1^{ml} \equiv \left(\frac{C_1^{ts}}{C_1^{ss}}\right)^2 \frac{\delta_s + C_2^{ss}}{\sigma_t^2 + \delta_{ts} + (C_2^{ts})^2 / C_2^{ss}}\,. \tag{104}$$

For this solution to be consistent with the condition $L_x^2 \gg L_h \gg L_x$, $N$ must satisfy $L_x^3 \gg B_1^{ml} N \gg L_x$ Therefore, if $B_1^{ml} N \gg L_x \gg (B_1^{ml} N)^{1/3}$, then the hidden layer size that minimizes the generalization error is

$$L_h^* \approx \sqrt{B_1^{ml} N L_x}\,. \tag{105}$$

###### Optimal hidden layer size when $L_x \gg L_h$

In this regime, the generalization error is given by

$$\epsilon_{gen} \approx \left(\sigma_t^2 + \delta_{ts} + \frac{(C_1^{ts})^2}{C_1^{ss}} + \frac{(C_2^{ts})^2}{C_2^{ss}} + \left(\frac{C_0^{ts}}{C_0^{ss}}\right)^2 \frac{\delta_s + C_1^{ss} + C_2^{ss}}{c_0 + L_h}\right) \frac{N}{N - L_h}\,. \tag{106}$$

Minimizing with respect to $L_h$ yields

$$L_h^* = \sqrt{(B_0^{ml})^2 + B_0^{ml}(N + c_0)} - B_0^{ml} - c_0 \tag{107}$$

where

$$B_0^{ml} \equiv \left(\frac{C_0^{ts}}{C_0^{ss}}\right)^2 \frac{\delta_s + C_1^{ss} + C_2^{ss}}{\sigma_t^2 + \delta_{ts} + (C_1^{ts})^2 / C_1^{ss} + (C_2^{ts})^2 / C_2^{ss}}\,. \tag{108}$$

For this solution to be consistent with the condition $L_x \gg L_h$, $N$ must satisfy $L_x \gg \sqrt{B_0^{ml} N}$. Assuming also that $N \gg 1$, we have

$$L_h^* \approx \sqrt{B_0^{ml} N}, \tag{109}$$

Here the optimal hidden layer size, $L_h^*$, does not depend on the input layer size, $L_x$.

#### 5.2 Stochastic Gradient Descent

For stochastic gradient descent, we use the cumulative generalization error, denoted $\epsilon_{cg}^N$ and defined to be

$$\epsilon_{cg}^N \equiv \frac{1}{N}\sum_{n=0}^{N-1}\left(\epsilon_{apr}+\sigma_t^2+\bar{\epsilon}_{est}^{(n)}\right) = \epsilon_{apr}+\sigma_t^2+\bar{\epsilon}_{cml}^N\,. \tag{110}$$

Using Eq. (89) for $\bar{\epsilon}_{cml}^N$ and Eq. (82) for $m_\infty^2$, from Eq. (77),

$$\epsilon_{cg}^N = (\epsilon_{apr}+\sigma_t^2)\left[2-\frac{1}{D_0^sL_h}\sum_q L_q\langle\lambda_\mu\rangle_q R_q(L_h)\right]+\sum_q R_q(L_h)\left[L_q\langle\lambda_\mu\rangle_q\frac{\sigma_R^2}{L_h}+\frac{(C_q^{ts})^2}{C_q^{ss}}\big(1-f_q(L_h)\big)\right]. \tag{111}$$

The critical quantity in this equation is $\langle\lambda_\mu\rangle_q$, which is given in Eq. (157) (but with slightly different notation). We repeat that equation here, following the notation used in §4.2, with a focus on the behavior when $L_h$ is either very small or very large,

$$\langle\lambda_\mu\rangle_0 \approx C_0^{ss}L_h \qquad L_h\gg 1 \tag{112a}$$

$$\langle\lambda_\mu\rangle_1 \approx \begin{cases} C_1^{ss}L_h/L_x & L_h\gg L_x\\ \delta_s+C_1^{ss}+C_2^{ss} & L_h\ll L_x\end{cases} \tag{112b}$$

$$\langle\lambda_\mu\rangle_2 \approx \begin{cases} 2C_2^{ss}L_h/L_x^2 & L_h\gg L_x^2\\ \delta_s+C_2^{ss}\left[1-L_x/L_h\right]^+ & L_h\ll L_x^2\end{cases} \tag{112c}$$

$$\langle\lambda_\mu\rangle_r = \delta_s\,. \tag{112d}$$

To make it easier to analyze the generalization error, it is convenient to use Eq. (88) to express $R_q(L_h)$ in terms of more fundamental quantities, yielding

$$\begin{aligned}\epsilon_{cg}^N \approx{}& (\epsilon_{apr}+\sigma_t^2)\left[2-\sum_{q\neq 0}\frac{L_q}{N}\left[1-e^{-N\langle\lambda_\mu\rangle_q/D_0^sL_h}\right]\right]+\frac{D_0^s}{N}\left[\sigma_R^2+\frac{(C_0^{ts})^2}{(C_0^{ss})^2}-\frac{\epsilon_{apr}+\sigma_t^2}{D_0^s}\right]\\ &+\sum_{q\neq 0}\frac{D_0^sL_q}{N}\left[1-e^{-N\langle\lambda_\mu\rangle_q/D_0^sL_h}\right]\left[\sigma_R^2+\frac{(C_q^{ts})^2}{C_q^{ss}}\frac{L_h}{L_q\langle\lambda_\mu\rangle_q}\big(1-f_q(L_h)\big)\right].\end{aligned} \tag{113}$$

To derive this expression, we replaced $\langle\lambda_\mu\rangle_0$ with $C_0^{ss}L_h$ (Eq. (112a)) and $f_0(L_h)$ with $c_0/L_h$ (Eq. (44a) in the large $L_h$ limit), used the fact that $L_0=1$ (Eq. (90b)), assumed $N\gg 1$, and replaced $(1-(C_0^{ss}/D_0^s))^N$ with $0$, which is valid in the large $N$ limit.

In the following subsections, we minimize $\epsilon_{cg}^N$ with respect to $L_h$ to find the optimal hidden layer size. As with maximum likelihood, we work in three different regimes, and again in each of them the estimation error becomes tractable. To simplify our analysis, though, we make assumptions about $N$ that assures the solution in each region is self-consistent.

###### Optimal hidden layer size when $L_h \gg L_x^2$

We assume that $N\gg L_x^2$, which will yield a self-consistent solution, as we show below. In the regime $L_h\gg L_x^2$, Eq. (35) tells us that $L_1=L_x, L_2\approx L_x^2/2$, and $L_r\approx L_h$. Consequently, using Eq. (88), with average eigenvalues given by Eq. (112), we see that $R_0(L_h)$, $R_1(L_h)$ and $R_2(L_h)$ are all approximately zero, and

$$R_r(L_h)=\frac{D_0^sL_h}{\delta_sN}\left(1-e^{-\frac{\delta_sN}{D_0^sL_h}}\right). \tag{114}$$

Inserting this into Eq. (111), using Eq. (112d) for $\langle\lambda_\mu^{(r)}\rangle$, recalling that $C_r^{ts}=0$ (see Eq. (90a)), and using the fact that all the other $R_q$ are approximately zero, we see that the cumulative generalization error is given approximately by

$$\epsilon_{cg}^N\approx\left(\epsilon_{apr}+\sigma_t^2\right)\left(2-\frac{L_h}{N}\left[1-e^{-\frac{\delta_sN}{D_0^sL_h}}\right]\right)+\frac{\sigma_R^2D_0^sL_h}{N}\left(1-e^{-\frac{\delta_sN}{D_0^sL_h}}\right). \tag{115}$$

The first term is a monotonically decreasing function of $L_h$ while the second term is monotonically increasing. When $\sigma_R^2$ is too small, the second term becomes too weak to supports the presence of the non-trivial minimum (gray points in

Fig. 5F). However, this initial weight dependence can be avoided by using an adaptive learning rate (black points in Fig. 5F), although the analytical estimation of the error becomes difficult in that case.

When $N \ll L_h$, the $L_h$ dependence in all but the term $\epsilon_{apr}$ in Eq. (115) disappears. We thus consider the opposite limit, $N \gg L_h$. Then, using Eq. (97) for the approximation error, we find, in this limit, that

$$\epsilon_{cg}^N \approx 2(\delta_{ts} + \sigma_t^2) + \left(\frac{C_2^{ts}}{C_2^{ss}}\right)^2 \frac{\delta_s L_x^2}{L_h} + \left(D_o^s \sigma_R^2 - [\delta_{ts} + \sigma_t^2]\right) \frac{L_h}{N} \, . \tag{116}$$

Minimizing with respect to $L_h$ gives

$$L_h^* = B_2^{sgd} \sqrt{N L_x^2}, \tag{117}$$

where

$$B_2^{sgd} \equiv \frac{C_2^{ts}}{C_2^{ss}} \sqrt{\frac{\delta_s}{D_0^s \sigma_R^2 - (\delta_{ts} + \sigma_t^2)}}. \tag{118}$$

We need to check for self-consistency, which means we need to check that $N \gg L_h^*$ and $L_h^* \gg L_x^2$. For the first, we combine the condition $L_x^2 \ll N$ with Eq. (117) to obtain $L_h^* \ll N$ (note that $B_2^{sgd}$ is $\mathcal{O}(1)$). For the second, we combine the condition $N \gg L_x^2$ with Eq. (117) to obtain $L_h^* \gg L_x^2$. Thus, this is a self-consistent solution.

##### Optimal hidden layer size when $L_x^2 \gg L_h \gg L_x$

In this regime we assume that $N \gg L_x$, which, as we show below, will again yield a self-consistent solution. Notably, this assumption is consistent with the assumed scaling in Fig. 5, $N \propto L_x^{1.9}$. In the regime $L_x^2 \gg L_h \gg L_x$, Eq. (35) tells us that $L_1 = L_x, L_2 \approx L_h$, and $L_r = 0$. Consequently, using Eq. (88), with average eigenvalues given by Eq. (112), we see that $R_0(L_h)$ and $R_1(L_h)$ are approximately zero, and

$$R_2(L_h) = \frac{D_0^s L_h}{N \langle \lambda_\mu^{(2)} \rangle} \left[1 - e^{-N \langle \lambda_\mu^{(2)} \rangle / (D_0^s L_h)}\right] \, . \tag{119}$$

Inserting this into Eq. (111), using Eq. (112c) for $\langle \lambda_\mu^{(2)} \rangle$, noting that $f_2(L_h) \approx f(2L_h/L_x^2; c_2) \approx 1$ (see Eq. (44c)), and using the fact that all the other $R_q$ are approximately zero, the cumulative generalization error is given approximately by

$$\epsilon_{cg}^N \approx (\epsilon_{apr} + \sigma_t^2) \left(2 - \frac{L_h}{N}\left[1 - e^{-\frac{(\delta_s + C_2^{ss})N}{D_o^s L_h}}\right]\right) + \frac{\sigma_R^2 D_o^s L_h}{N}\left[1 - e^{-\frac{(\delta_s + C_2^{ss})N}{D_0^s L_h}}\right] \, . \tag{120}$$

Following the arguments in the previous section, we consider the limit $N \gg L_h$. Then, using Eq. (97) for the approximation error, we find, in this limit, that

$$\epsilon_{cg}^N \approx 2\left(\delta_{ts} + \sigma_t^2 + \frac{(C_2^{ts})^2}{C_2^{ss}}\right) + 2\left(\frac{C_1^{ts}}{C_1^{ss}}\right)^2 \frac{(\delta_s + C_2^{ss}) L_x}{L_h} + \left(D_o^s \sigma_R^2 - \left[\delta_{ts} + \sigma_t^2 + \frac{(C_2^{ts})^2}{C_2^{ss}}\right]\right) \frac{L_h}{N} \, . \tag{121}$$

Minimizing with respect to $L_h$ yields

$$L_h^* = B_1^{sgd} \sqrt{N L_x} \, , \tag{122}$$

where

$$B_1^{sgd} \equiv \frac{C_1^{ts}}{C_1^{ss}} \sqrt{\frac{2(\delta_s + C_2^{ss})}{D_o^s \sigma_R^2 - [\delta_{ts} + (C_2^{ts})^2 / C_2^{ss} + \sigma_t^2]}} \, . \tag{123}$$

We need to check for self-consistency, which means we need to check that $N \gg L_h^*$ and $L_x^2 \gg L_h^* \gg L_x$. For the first, we combine the condition $L_x \ll N$ with Eq. (122) to obtain $L_h^* \ll N$ (note that $B_1^{sgd}$ is $\mathcal{O}(1)$). For the second, we combine the condition $N \gg L_x$ with Eq. (122) to obtain $L_h^* \gg L_x$. Thus, this is a self-consistent solution. To ensure $L_x^2 \gg L_h^*$, $N$ needs to satisfy $N \ll L_x^3$. Thus, $N$ must be large but not too large.

##### Optimal hidden layer size when $L_x \gg L_h$

In this regime, we assume that $N \gg 1$, which is again consistent with the assumed scaling in Fig. 5, $N \propto L_x^{1.9}$. In the regime $L_x \gg L_h$, Eq. (35) tells us that $L_1 = L_h$ and $L_2 = L_r = 0$. Because $L_2$ and $L_r$ are zero and $N \gg 1$, $q = 1$ is the only relevant term, so cumulative generalization error, Eq. (111), is given approximately by

$$\epsilon_{cg}^N \approx (\epsilon_{apr} + \sigma_t^2)\left(2 - \frac{L_h}{N}\left[1 - e^{-\frac{\langle\lambda^{(1)}\rangle N}{D_0^s L_h}}\right]\right) + \frac{\sigma_R^2 D_0^s L_h}{N}\left(1 - e^{-\frac{\langle\lambda^{(1)}\rangle N}{D_0^s L_h}}\right). \tag{124}$$

As before, using Eq. (97) and assuming $N \gg L_h$, the above equation becomes

$$\epsilon_{cg}^N \approx 2\left(\frac{C_0^{ts}}{C_0^{ss}}\right)^2 \frac{\delta_s + C_1^{ss} + C_2^{ss}}{L_h} + \left(D_0^s\sigma_R^2 - \left[\sigma_t^2 + \delta_{ts} + \frac{(C_1^{ts})^2}{C_1^{ss}} + \frac{(C_2^{ts})^2}{C_2^{ss}}\right]\right)\frac{L_h}{N} + \text{const.} \tag{125}$$

Thus, the optimal hidden layer size follows

$$L_h^* = B_0^{sgd}\sqrt{N}, \tag{126}$$

where

$$B_0^{sgd} \equiv \frac{C_0^{ts}}{C_0^{ss}}\sqrt{\frac{2(\delta_s + C_1^{ss} + C_2^{ss})}{D_0^s\sigma_R^2 - [\delta_{ts} + (C_1^{ts})^2/C_1^{ss} + (C_2^{ts})^2/C_2^{ss} + \sigma_t^2]}}. \tag{127}$$

This is a self-consistent solution at $L_x^2 \gg N$.

### 6 Model with low precision hard-wired connections

We extend the student model by adding a parallel hidden layer corresponding to lateral horn neurons,

$$y = \boldsymbol{w}_p \cdot g(\boldsymbol{J}_p\boldsymbol{x}) + \boldsymbol{w}_s \cdot g(\boldsymbol{J}_s\boldsymbol{x}). \tag{128}$$

Here, $g$ is ReLU, $\boldsymbol{J}_p \in \Re^{L_p \times L_x}$, and $\boldsymbol{w}_p \in \Re^{L_p}$. We hypothesize that $\boldsymbol{J}_p^*$ and $\boldsymbol{w}_p^*$ are genetically encoded, and these weights are optimized on evolutionary timescales. Thus, ideally, $\boldsymbol{J}_p$ and $\boldsymbol{w}_p$ should be chosen as

$$\boldsymbol{J}_p^*, \boldsymbol{w}_p^* = \arg\min_{\boldsymbol{J}_p, \boldsymbol{w}_p} \left\langle \left[\boldsymbol{w}_t \cdot g(\boldsymbol{J}_t\boldsymbol{x}) - \boldsymbol{w}_p \cdot g(\boldsymbol{J}_p\boldsymbol{x})\right]^2 \right\rangle_{p(\boldsymbol{x})}. \tag{129}$$

However, this optimization is not analytically tractable unless $L_p > L_t$, and, more importantly, encoding $\boldsymbol{J}_p^*$ and $\boldsymbol{w}_p^*$ perfectly requires an infinite number of bits, while genetic capacity is limited. Thus, to genetically hard-wire the lateral horn pathway, the weights need to be compressed. Below, we first describe the greedy optimization method we used for estimating $\boldsymbol{J}_p^*$ and $\boldsymbol{w}_p^*$, then explain two methods for compressing the weights.

##### Numerical estimation of $\boldsymbol{J}_p^*$ and $\boldsymbol{w}_p^*$

We used a mini-batch backpropagation for updating $\boldsymbol{J}_p$,

$$\boldsymbol{J}_p^{(m+1)} = \boldsymbol{J}_p^{(m)} - \eta\sum_{b=1}^{B}\frac{\partial}{\partial \boldsymbol{J}_p}\left(\boldsymbol{w}_p \cdot g(\boldsymbol{J}_p\boldsymbol{x}_b) - \boldsymbol{w}_t \cdot g(\boldsymbol{J}_t\boldsymbol{x}_b)\right)^2, \tag{130}$$

where $m$ is the update count. We used mini-batch size $B = 500$, learning rate $\eta = 0.02$, and the teacher noise was excluded from the supervised signal to achieve fast convergence. Because the optimization of $\boldsymbol{J}_p$ is an evolutionary process, the update rule does not need to be local. The weight, $\boldsymbol{w}_p$, was updated after each minibatch update of $\boldsymbol{J}_p$ according to

$$\boldsymbol{w}_p^{(m)} \equiv \left\langle g(\boldsymbol{J}_p^{(m)}\boldsymbol{x})g(\boldsymbol{J}_p^{(m)}\boldsymbol{x})^T\right\rangle^{-1}\left\langle g(\boldsymbol{J}_p^{(m)}\boldsymbol{x})g(\boldsymbol{J}_t\boldsymbol{x})^T\right\rangle \boldsymbol{w}_t. \tag{131}$$

The above expectations, which are over $\boldsymbol{x}$, were obtained analytically using Eq. (144) below. We initialized $\boldsymbol{J}_p^{(m=0)}$ to $J_{ij}^{p,0} \sim N(0, 1/L_x)$, then updated $\boldsymbol{w}_p$ and $\boldsymbol{J}_p$ alternatively for $10^5$ steps (see §7.4 for an algorithmic description), which was typically enough to achieve convergence.

###### Compression of the weights by discretization

Synaptic weights can be compressed by discretization or by adding noise [25]. Although, we mainly used the latter, we describe the discretization-based approach first, as it is more intuitive.

Real-valued weights, $w_j^*$, can be compressed to $s_b$ bits by simply discretizing them into $2^{s_b}$ states. Denoting $w_{max} \equiv \max\{w_1^*, ..., w_{L_p}^*\}$, $w_{min} \equiv \min\{w_1^*, ..., w_{L_p}^*\}$, and $\Delta w \equiv (w_{max} - w_{min})/(2^{s_b})$, compressed weights, $w_j$, are obtained via

$$w_j = w_{min} + \left( \left\lfloor \frac{w_j^* - w_{min}}{\Delta w} \right\rfloor + \frac{1}{2} \right) \Delta w, \tag{132}$$

where $\lfloor x \rfloor$ returns the largest integer less than or equal to $x$. Both the hidden and output weights, $\boldsymbol{J}_p$ and $\boldsymbol{w}_p$, can be compressed in this manner.

###### Compression of the weights by adding noise

Assuming that the optimal weight $w_j^*$ is sampled from a Gaussian distribution (i.e., $w_j^* \sim N(0, \sigma_w^2)$), the bit length required for encoding the weight can be reduced by shrinking the weight while adding noise,

$$w_j = \sqrt{1-\gamma^2} w_j^* + \gamma \sigma_w \zeta, \tag{133}$$

where $\zeta$ is a zero mean, unit variance Gaussian variable, and $\gamma$ is the relative noise amplitude ($0 \leq \gamma \leq 1$). Marginalizing over $w_j^*$, we get $w_j \sim N(0, \sigma_w^2)$. Thus, the mutual information between $w_j$ and $w_j^*$ is

$$I[w_j; w_j^*] = -\log \gamma. \tag{134}$$

Therefore, to compress the weight into $s_b$ bits, $\gamma$ needs to be set to $\gamma = e^{-(\log 2) s_b}$ ($\log 2$ is for the conversion from nats to bits). Note that in this formulation, $2^{s_b}$ does not need to be an integer, unlike for the discretization method.

In the simulations, we estimated the variance of $\{w_j^*\}$ and $\{J_{ij}^*\}$ numerically, after they were optimized. Denoting the variances as $\sigma_w^2$ and $\sigma_J^2$, the compressed weights are given as

$$w_j = \sqrt{1-\gamma^2} w_j^* + \gamma \sigma_w \zeta \quad \text{and} \quad J_{ij} = \sqrt{1-\gamma^2} J_{ij}^* + \gamma \sigma_J \zeta. \tag{135}$$

### 7 Details of numerical analysis

#### 7.1 ReLU activation

We first compute the parameters when both the teacher and student use ReLU activation ($g_t(u) = g_s(u) = \max(0, u)$). The diagonal elements, Eqs. (15) and (37), are given by

$$D_0^s = D_0^t = \int_0^\infty \frac{1}{\sqrt{2\pi}} \exp\left(-\frac{u^2}{2}\right) du = \frac{1}{2}\,. \tag{136}$$

The off diagonal elements, Eqs. (19) and (31), are given by

$$C_0^{ts} = C_0^{ss} = \frac{1}{2\pi} \tag{137a}$$

$$C_1^{ts} = C_1^{ss} = \frac{1}{4} \tag{137b}$$

$$C_2^{ts} = C_2^{ss} = \frac{1}{4\pi}\,. \tag{137c}$$

In this setting, the coefficient for the third order term is identically zero, so the second-order approximation effectively achieves third-order accuracy. Inserting the coefficients into Eq. (43), the approximation error is estimated as

$$\begin{aligned} \epsilon_{apr} \approx\; & \delta_s + \frac{1}{4\pi}\left(1 - \left[1 - \frac{L_x}{L_h}\right]^+\right) + \frac{\pi - 1}{2\pi(\pi - 1 + L_h)} \\ & + \frac{1}{8}\left[\sqrt{\left(\frac{L_h}{L_x} + 4\delta_s + \frac{1}{\pi} - 1\right)^2 + 4\left(4\delta_s + \frac{1}{\pi}\right)} - \left(\frac{L_h}{L_x} + 4\delta_s + \frac{1}{\pi} - 1\right)\right] \\ & + \frac{1}{8\pi}\left[1 - \frac{L_x}{L_h}\right]^+ \left[\sqrt{\left(\frac{2L_h}{L_x^2} + \frac{4\pi\delta_s}{1 - L_x/L_h} - 1\right)^2 + \frac{16\pi\delta_s}{1 - L_x/L_h}} - \left(\frac{2L_h}{L_x^2} + \frac{4\pi\delta_s}{1 - L_x/L_h} - 1\right)\right] \end{aligned} \tag{138}$$

where $\delta_s = (\pi - 3)/4\pi$ (see Eq. (24)).

#### 7.2 Logistic activation

We next consider the case of model mismatch, where the teacher activation function is ReLU but the student is a logistic function, $g_s(u) = 1/(1+e^{-u})$. The diagonal element of the teacher, $D_0^t$, is the same as above, but the student is different,

$$D_0^s = \int_{-\infty}^{\infty} \frac{du}{\sqrt{2\pi}} \left( \frac{1}{1+e^{-u}} \right)^2 \exp\left( -\frac{u^2}{2} \right) \simeq 0.29338 \,. \tag{139}$$

The diagonal elements are given by

$$C_0^{ss} = \frac{1}{4} \tag{140a}$$
$$C_1^{ss} \simeq 0.04269 \tag{140b}$$
$$C_2^{ss} = 0 \tag{140c}$$
$$C_0^{ts} = \frac{1}{2\sqrt{2\pi}} \tag{140d}$$
$$C_1^{ts} \simeq 0.1033 \tag{140e}$$
$$C_2^{ts} = 0. \tag{140f}$$

Here, $\simeq$ represents a numerical approximation of an integral. Notably, because both $C_2^{ss}$ and $C_2^{ts}$ are zero, the second-order term disappears from the approximation error. Thus, using $c_0 = (\delta_s + C_1^{ss})/C_0^{ss}$, the approximation error simplifies to

$$\epsilon_{apr} \approx \left( \frac{1}{2} - \frac{(C_0^{ts})^2}{C_0^{ss}} - \frac{(C_1^{ts})^2}{C_1^{ss}} \right) + \frac{(C_0^{ts})^2}{C_0^{ss}} \frac{c_0}{c_0 + L_h} + \frac{(C_1^{ts})^2}{2C_1^{ss}} \left( \sqrt{\left[ \frac{L_h}{L_x} + \frac{\delta_s}{C_1^{ss}} - 1 \right]^2 + \frac{4\delta_s}{C_1^{ss}}} - \left[ \frac{L_h}{L_x} + \frac{\delta_s}{C_1^{ss}} - 1 \right] \right) . \tag{141}$$

#### 7.3 Numerical estimation of the errors

The generalization error is easily estimated numerically by evaluating the test error over a large number of test samples,

$$\epsilon_{gen} \approx \frac{1}{N_{test}} \sum_{n=1}^{N_{test}} \left( \boldsymbol{w}_s \cdot g(\boldsymbol{J}_s \boldsymbol{x}_n) - y_n \right)^2 . \tag{142}$$

In simulations of maximum likelihood learning, we calculated the weights using Eq. (49), then computed $\epsilon_{gen}$ using $N_{test} = 30,000$ samples. The cumulative generalization error under SGD learning was estimated using

$$\epsilon_{cg}^N \approx \frac{1}{N} \sum_{n=1}^{N} \left( \boldsymbol{w}_s^{(n-1)} \cdot g(\boldsymbol{J}_s \boldsymbol{x}_n) - y_n \right)^2 . \tag{143}$$

Note that, because we provided a new sample $\{\boldsymbol{x}_n, y_n\}$ in each update, the right hand side is the cumulative test error, not the training error.

Estimating the approximation error (Eq. (10)), and the estimation error (Eq. (47)) from simulations is harder, becuase we need to evaluate $\langle \boldsymbol{g}_t \boldsymbol{g}_t^T \rangle$, $\langle \boldsymbol{g}_t \boldsymbol{g}_s^T \rangle$, and $\langle \boldsymbol{g}_s \boldsymbol{g}_s^T \rangle$ and the averages over $\boldsymbol{x}$ are generally intractable due to the high-dimensionality. However, if the nonlinearity, $g(\cdot)$, in both the teacher and student networks are ReLU, marginalization over $\boldsymbol{u} \equiv \boldsymbol{J}\boldsymbol{x}$ has a closed-form expression,

$$\begin{aligned} \langle g_q(u_i) g_{q'}(u_j) \rangle_{p(u_i,u_j)} &= \int_0^{\infty} du_i \int_0^{\infty} du_j \frac{u_i u_j}{2\pi\sigma_i\sigma_j\sqrt{1-\rho_{ij}^2}} \exp\left( -\frac{1}{2(1-\rho_{ij}^2)} \left[ \frac{u_i^2}{\sigma_i^2} + \frac{u_j^2}{\sigma_j^2} - \frac{2\rho_{ij}u_i u_j}{\sigma_i\sigma_j} \right] \right) \\ &= \frac{\sigma_i\sigma_j}{2\pi} \left( \sqrt{1-(\rho_{ij})^2} + \rho_{ij}\cos^{-1}(-\rho_{ij}) \right) \end{aligned} \tag{144}$$

where $\sigma_i^2 = (\boldsymbol{J}_q \boldsymbol{J}_q^T)_{ii}$, $\rho_{ij} = \frac{(\boldsymbol{J}_q \boldsymbol{J}_{q'}^T)_{ij}}{\sigma_i\sigma_j}$, and the indices $q$ and $q'$ are either $s$ or $t$. The errors under a specific network realization in Figs. 3, 5B, and 5C were calculated using this expression. For numerical stability, the inverse, $\boldsymbol{G}_s^{-1}$, was computed by solving a linear matrix equation $\boldsymbol{G}_s \boldsymbol{w}_s = \langle \boldsymbol{g}_s \boldsymbol{g}_t^T \rangle \boldsymbol{w}_t$, (see Eq. (6) and *github:nhiratani/olfactory_design*).

For the logistic activation, we only computed (and plotted) the generalization error, as numerical estimation of the approximation/estimation errors is difficult in this setting.

#### 7.4 Model with evolutionary and developmental learning

In the model with low-precision hard-wired connections, the learning process is described as follows,

Initialize $\boldsymbol{J}_p^*$ by $J_{ij}^{P*} \sim N(0, 1/L_x)$;
**for** $m = 1, ..., 10^5$ **do**
  Update $\boldsymbol{w}_p^*$ and $\boldsymbol{J}_p^*$ using Eq. (131) and Eq. (130), alternatively.
**end**
Compress $\boldsymbol{w}_p^*$ and $\boldsymbol{J}_p^*$ into $s_b$ bits weights $\boldsymbol{w}_p$ and $\boldsymbol{J}_p$ using Eq. (135);
Initialize $\boldsymbol{J}_s$ and $\boldsymbol{w}_s$ by $\boldsymbol{J}_s \sim N(0, 1/L_x)$ and $\boldsymbol{w}_s = \boldsymbol{0}$;
**for** $n = 1, ..., N$ **do**
  $\boldsymbol{x}_n \sim N(0, I)$, $y_n \sim N(\boldsymbol{w}_t \cdot g(\boldsymbol{J}_t \boldsymbol{x}_n), \sigma_t^2)$;
  $\boldsymbol{w}_s^{(n)} = \boldsymbol{w}_s^{(n-1)} + \frac{2}{\max(L_h, n)}(y_n - [\boldsymbol{w}_p \cdot \boldsymbol{g}(\boldsymbol{J}_p \boldsymbol{x}_n) + \boldsymbol{w}_s^{(n-1)} \cdot g(\boldsymbol{J}_s \boldsymbol{x}_n)])g(\boldsymbol{J}_s \boldsymbol{x}_n)$;
**end**

In Fig. 6C we instead used Eq. (132) for the compression of the weights $\boldsymbol{J}_p^*$ and $\boldsymbol{w}_p^*$. In Fig. 6D, the compression was done with Eq. (135), and $\boldsymbol{w}_p$ was additionally trained in the developmental learning phase using

$$\boldsymbol{w}_p^{(n)} = \boldsymbol{w}_p^{(n-1)} + \frac{2}{\max(L_h, n)}(y_n - [\boldsymbol{w}_p^{(n-1)} \cdot g(\boldsymbol{J}_p \boldsymbol{x}_n) + \boldsymbol{w}_s^{(n-1)} \cdot g(\boldsymbol{J}_s \boldsymbol{x}_n)])g(\boldsymbol{J}_p \boldsymbol{x}_n) \tag{145}$$

from the low-precision weight $\boldsymbol{w}_p^{(n=0)}$ derived from Eq. (135).

#### 7.5 Numerical estimation of the optimal hidden layer size

In both maximum likelihood and SGD simulations, we first estimated the generalization error by calculating the mean error over $K_{\text{sim}}$ simulations, for various $L_h$ spanning from $L_h = 10$ to $L_h = L_h^{\max}$ with 10% increment at each step. Then, we defined the empirical estimate of the optimal hidden layer size as the network size that yielded the minimum average error.

In the maximum likelihood simulations, we used $L_h^{\max} = \min[N, 30,000]$, except for the large $L_x$ simulations in Fig. 4A, where we used $L_h^{\max} = 15,000$ for $L_x > 10,000$, $L_h^{\max} = 6,000$ for $L_x > 30,000$, and in Fig. 4E, where we set $L_h^{\max} = 4,000$. For each $L_x$, we took the mean over $K_{\text{sim}} = 100$ simulations if $N < 1000$, else $K_{\text{sim}} = 10$ simulations.

In the SGD simulations, we set $L_h^{\max} = 30,000$ and $K_{\text{sim}} = 10$ simulations, except for Fig. 5B where we used $K_{\text{sim}} = 100$, and for the large $L_x$ region of Fig. 4D, where we set $L_h^{\max} = 10,000$ for $L_x > 7,000$, and $L_h^{\max} = 3,300$ for $L_x > 20,000$). In Fig. 5F and Fig. 6, we used $L_h^{\max} = 100,000$.

### 8 Eigenvectors and eigenvalues of $\boldsymbol{G}_s$

Here we estimate the eigenvectors and eigenvalues of $\boldsymbol{G}_s$, based on the approximate expression given in Eq. (28). That expression consists of four matrices: the identity, a rank one matrix with eigenvalue that scales as $L_h$, and, as pointed out immediately after Eq. (28), two matrices with Marchenko-Pastur distributions for their eigenvalues,

$$\boldsymbol{J}_s \boldsymbol{J}_s^T\colon\ \lambda \sim MP(1, L_h/L_x) \tag{146a}$$

$$\boldsymbol{M}_s \boldsymbol{M}_s^T\colon\ \lambda \sim MP(1, 2L_h/L_x^2)\,. \tag{146b}$$

For these matrices, so long as $L_h > L_x^2/2$, the nonzero eigenvalues scale as $L_h/L_x$ and $2L_h/L_x^2$, respectively. In this regime, the nonzero eigenvalues of the three non-identity matrices in Eq. (28) are successively smaller, each time by a factor of $L_x$. We will assume this holds in general; when it does not, our approximation may not be very accurate.

To make use of the successively smaller eigenvalues, we note that if we sum two matrices with very different eigenvalues, the one with large eigenvalues dominates. More formally, consider two symmetric matrices, $\boldsymbol{Q}$ and $\boldsymbol{R}$, such that their nonzero eigenvalues are both $\mathcal{O}(1)$. Letting $\boldsymbol{v}_Q$ be an eigenvector of $\boldsymbol{Q}$ with eigenvalue $\lambda_Q$, for $|\epsilon| \ll 1$, we have

$$(\boldsymbol{Q} + \epsilon \boldsymbol{R})\boldsymbol{v}_Q = \boldsymbol{Q}\boldsymbol{v}_Q + \epsilon \boldsymbol{R}\boldsymbol{v}_Q = \lambda_Q \boldsymbol{v}_Q + \epsilon \boldsymbol{R}\boldsymbol{v}_Q \approx \lambda_Q \boldsymbol{v}_Q\,. \tag{147}$$

If $\boldsymbol{Q}$ is rank-deficient, there will be additional $\mathcal{O}(\epsilon)$ eigenvalues. Their eigenvectors will lie in the space spanned by $\boldsymbol{R}$, but with the space spanned by $\boldsymbol{Q}$ projected out.

We will now apply this to $\boldsymbol{G}_s$, but with a small correction, which turns out to be necessary to get good agreement with simulations: when computing the eigenvalue associated with the rank one matrix $\boldsymbol{1}_h \boldsymbol{1}_h^T$, we treat $\boldsymbol{J}_s \boldsymbol{J}_s^T$ and $\boldsymbol{M}_s \boldsymbol{M}_s^T$ as identity matrices, and when computing the eigenvalue spectrum associated with $\boldsymbol{J}_s \boldsymbol{J}_s^T$ we treat $\boldsymbol{M}_s \boldsymbol{M}_s^T$ as the identity matrix. (Note that $\boldsymbol{J}_s\boldsymbol{J}_s^T$ and $\boldsymbol{M}_s\boldsymbol{M}_s^T$ are typically rank deficient. However, we can consider an ensemble average; because their eigenvalues average to 1, that ensemble average is the identity matrix.)

Using this procedure, the relevant eigenvalue equation associated with the matrix associated with $\mathbf{1}_h\mathbf{1}_h^T$ is

$$\left((\delta_s + C_1^{ss} + C_2^{ss})\boldsymbol{I} + C_0^{ss}\mathbf{1}_h\mathbf{1}_h^T\right)\cdot \boldsymbol{v}^{(0)} = \lambda^{(0)}\boldsymbol{v}^{(0)}\,, \tag{148}$$

implying that

$$\lambda^{(0)} = \delta_s + C_1^{ss} + C_2^{ss} + C_0^{ss}L_h \tag{149a}$$

$$\boldsymbol{v}^{(0)} = \frac{\mathbf{1}_h}{\sqrt{L_h}}\,. \tag{149b}$$

To find the eigenvalues associated with $\boldsymbol{J}_s\boldsymbol{J}_s^T$, we should project out the one dimensional subspace spanned by $\mathbf{1}_h$, but that will have an $\mathcal{O}(1/L_h)$ effect, so we do not do it. Consequently, the relevant eigenvalue equation associated with the matrix $\boldsymbol{J}_s\boldsymbol{J}_s^T$ is

$$\left((\delta_s + C_2^{ss})\boldsymbol{I} + C_1^{ss}\boldsymbol{J}_s\boldsymbol{J}_s^T\right)\boldsymbol{v}_k^{(1)} = \lambda_k^{(1)}\boldsymbol{v}_k^{(1)}\,, \tag{150}$$

implying that

$$\lambda_k^{(1)} = \delta_s + C_2^{ss} + C_1^{ss}\tilde{\lambda}_k^{(1)} \tag{151}$$

where

$$\tilde{\lambda}^{(1)} \sim MP^+\left(1, \frac{L_h}{L_x}\right)\,. \tag{152}$$

The $+$ superscript on $MP$ indicates that we should include only the non-zero eigenvalues.

To find the eigenvalues associated with $\boldsymbol{M}_s\boldsymbol{M}_s^T$, we need to project out the subsapce spanned by $\boldsymbol{J}_s\boldsymbol{J}_s^T$. Using $\widetilde{\boldsymbol{M}}_s$ to denote $\boldsymbol{M}_s$ in the lower dimensional space, the relevant eigenvalue equation is

$$\left(\delta_s\boldsymbol{I} + C_2^{ss}\widetilde{\boldsymbol{M}}_s\widetilde{\boldsymbol{M}}_s^T\right)\boldsymbol{v}_k^{(2)} = \lambda_k^{(2)}\boldsymbol{v}_k^{(2)}\,, \tag{153}$$

The dimension of the subspace we project out is $\max[L_h, L_x]$. Assuming that the projection $\boldsymbol{M}\to\widetilde{\boldsymbol{M}}$ is random, the eigenvalues of $\boldsymbol{M}_s\boldsymbol{M}_s^T$ are reduced by a factor of $[1 - L_x/L_h]^+$, giving us

$$\lambda_k^{(2)} = \delta_s + C_2^{ss}\tilde{\lambda}_k^{(2)} \tag{154}$$

where

$$\tilde{\lambda}^{(2)} \sim MP^+\left(\left[1 - \frac{L_x}{L_h}\right]^+, \frac{2L_h}{L_x^2}\right)\,. \tag{155}$$

Finally, if $L_r > 0$, there are additional eigenvectors. We have already taken care of the matrices with structure, so the remaining matrix is just $\delta_s\boldsymbol{I}$. Consequantly,

$$\lambda_k^{(r)} = \delta_s\,. \tag{156}$$

Because we need them for the analysis of SGD, we compute the average eigenvalues for the two components: $\tilde{\lambda}^{(1)}$ and $\tilde{\lambda}^{(2)}$. For the full Marchenko-Pastur distribution with parameters $\sigma^2$ and $\lambda$, the average eigenvalue is $\sigma^2$. However, for the distribution over only the non-zero eigenvalues, the average eigenvalue is $\sigma^2\max[1,\lambda]$. Thus,

$$\lambda^{(0)} = \delta_s + C_1^{ss} + C_2^{ss} + C_0^{ss}L_h \tag{157a}$$

$$\langle\lambda_k^{(1)}\rangle = \delta_s + C_2^{ss} + C_1^{ss}\max\left[1, \frac{L_h}{L_x}\right] \tag{157b}$$

$$\langle\lambda_k^{(2)}\rangle = \delta_s + C_2^{ss}\left[1 - \frac{L_x}{L_h}\right]^+\max\left[1, \frac{2L_h}{L_x^2}\right] \tag{157c}$$

$$\langle\lambda_k^{(r)}\rangle = \delta_s \tag{157d}$$

where we included $\lambda^{(0)}$(Eq. (149a)), and $\lambda_r$ for completeness.

To compute the approximation error, we also need the eigenvalue/eigenvector expansion of the right hand side of Eq. (39). Repeating the above analysis, we find that

$$\lambda_k^{ts(0)} = \frac{(C_1^{ts})^2}{L_x} + \frac{(C_2^{ts})^2}{L_x^2/2} + (C_0^{ts})^2 L_h \approx (C_0^{ts})^2 L_h \tag{158a}$$

$$\lambda_k^{ts(1)} = \frac{(C_2^{ts})^2}{L_x^2/2} + \frac{(C_1^{ts})^2}{L_x}\tilde{\lambda}_k^{(1)} \approx \frac{(C_1^{ts})^2}{L_x}\tilde{\lambda}_k^{(1)} \tag{158b}$$

$$\lambda_k^{ts(2)} = \frac{(C_2^{ts})^2}{L_x^2/2}\tilde{\lambda}_k^{(2)} \tag{158c}$$

$$\lambda_k^{ts(r)} = 0 \tag{158d}$$

where the approximations are valid in the large $L_x$ limit.

### 9 Marchenko-Pastur averages

In §3 (see in particular Eq. (40)) we need to compute averages of the form

$$\frac{1}{L}\sum_{k=1}^{L'} \frac{\lambda_k}{c+\lambda_k} = \frac{L'}{L}\left\langle \frac{\lambda}{c+\lambda} \right\rangle_{\lambda\sim MP^+(\sigma^2,\bar{\lambda})} \tag{159}$$

where, as above the $+$ superscript on $MP$ indicates that the average is over only the positive eigenvalues. Computing analytically the average on the right hand side, we have

$$\frac{L'}{L}\left\langle \frac{\lambda}{c+\lambda} \right\rangle_{\lambda\sim MP^+(\sigma^2,\bar{\lambda})} = \frac{L'/L}{\min\left[1,\bar{\lambda}\right]}\left(1 - f\left(\bar{\lambda};\frac{c}{\sigma^2}\right)\right) \tag{160}$$

where

$$f(\bar{\lambda};c) \equiv \frac{\sqrt{\left(\bar{\lambda}-1+c\right)^2 + 4c} - \left(\bar{\lambda}-1+c\right)}{2}\,. \tag{161}$$

The large and small $\bar{\lambda}$ limits of $f$ are relatively simple,

$$f(\bar{\lambda}, c/\sigma^2) \to \begin{cases} 1 & \bar{\lambda}\to 0 \\ c/(\sigma^2\bar{\lambda}) & \bar{\lambda}\to\infty\,. \end{cases} \tag{162}$$

The small $\bar{\lambda}$ limit is important, because it tells us that $1 - f(\bar{\lambda}, c/\sigma^2)$ is small whenever $\bar{\lambda}$ is small.

For the first sum in Eq. (40), $L = L_x$, $L' = L_1 = \min[L_x, L_h]$, and $c = c_1$ (defined in Eq. (34b)). The parameters of the Marchenko-Pastur distribution, given in Eq. (152), are $\sigma^2 = 1$ and $\bar{\lambda} = L_h/L_x$. The latter implies that $L'/(L\min[1,\bar{\lambda}]) = 1$. Consequently, the first sum in Eq. (40) is

$$\frac{1}{L_x}\sum_{k=1}^{L_1} \frac{\tilde{\lambda}_k^{(1)}}{c_1+\tilde{\lambda}_k^{(1)}} = 1 - f(\bar{\lambda}; c_1)\,. \tag{163}$$

For the second sum in Eq. (40), $L = L_x^2/2$, $L' = L_2 = \min[L_x^2/2, L_h - L_x]^+$ and $c = c_2$ (defined in Eq. (34c). The parameters of the Marchenko-Pastur distribution, given in Eq. (155), are $\sigma^2 = [1 - L_x/L_h]^+$ and $\bar{\lambda} = 2L_h/L_x^2$. We thus have, after a small amount of algebra,

$$\frac{1}{L_x^2/2}\sum_{k=1}^{L_2} \frac{\tilde{\lambda}_k^{(2)}}{c_2+\tilde{\lambda}_k^{(2)}} = \frac{\min[L_x^2/2, L_h - L_x]^+}{\min[L_x^2/2, L_h]^+}\left(1 - f\left(\frac{L_h}{L_x^2/2}; \frac{c_2}{[1-L_x/L_h]^+}\right)\right)\,. \tag{164}$$

Noticing that the first term is 1 at $L_h > L_x^2/2 + L_x$, $[1 - L_x/L_h]^+$ at $L_h < L_x^2/2$, and slightly smaller than 1 in between, we can simplify the expression above as

$$\frac{1}{L_x^2/2}\sum_{k=1}^{L_2} \frac{\tilde{\lambda}_k^{(2)}}{c_2+\tilde{\lambda}_k^{(2)}} \approx \left[1 - \frac{L_x}{L_h}\right]^+ \left[1 - f\left(\frac{L_h}{L_x^2/2}; \frac{c_2}{1-L_x/L_h}\right)\right]\,. \tag{165}$$

In §4.1 we need the average of the inverse of the eigenvalue. That is easily found from the above analysis,

$$\left\langle \frac{1}{\lambda} \right\rangle_{\lambda\sim MP^{+}(\sigma^2,\bar{\lambda})} = \frac{\partial}{\partial c}\bigg|_{c=0} \left\langle \frac{-\lambda}{c+\lambda} \right\rangle_{\lambda\sim MP^{+}(\sigma^2,\bar{\lambda})} . \tag{166}$$

Using Eq. (160) for the right hand side, a straightforward calculation yields

$$\left\langle \frac{1}{\lambda} \right\rangle_{\lambda\sim MP^{+}(\sigma^2,\bar{\lambda})} = \frac{1}{\sigma^2|\bar{\lambda}-1|} . \tag{167}$$

[21] Robi Tacutu, Thomas Craig, Arie Budovsky, Daniel Wuttke, Gilad Lehmann, Dmitri Taranukha, Joana Costa, Vadim E Fraifeld, and João Pedro De Magalhães. Human ageing genomic resources: integrated databases and tools for the biology and genetics of ageing. *Nucleic acids research*, 41(D1):D1027–D1033, 2012.

[22] Shyam Srinivasan and Charles F Stevens. Scaling principles of distributed circuits. *Current Biology*, 29(15):2533–2540, 2019.

[23] Vladimir A Mar$vc$enko and Leonid Andreevich Pastur. Distribution of eigenvalues for some sets of random matrices. *Mathematics of the USSR-Sbornik*, 1(4):457, 1967.

[24] Madhu Advani and Surya Ganguli. Statistical mechanics of optimal convex inference in high dimensions. *Physical Review X*, 6(3):031034, 2016.

[25] Geoffrey Hinton and Drew Van Camp. Keeping neural networks simple by minimizing the description length of the weights. In *in Proc. of the 6th Ann. ACM Conf. on Computational Learning Theory*. Citeseer, 1993.
